# High performance intensiometric direct- and inverse-response genetically encoded biosensors for citrate

**DOI:** 10.1101/2020.04.12.038547

**Authors:** Yufeng Zhao, Yi Shen, Yurong Wen, Robert E. Campbell

**Author notes:** Correspondence: Regarding x-ray crystallography should be addressed to Y.W. All other correspondence should be addressed to R.E.C.

## Abstract

Motivated by the growing recognition of citrate as a central metabolite in a variety of biological processes associated with healthy and diseased cellular states, we have developed a series of high-performance genetically encoded citrate biosensors suitable for imaging of citrate concentrations in mammalian cells. The design of these biosensors was guided by structural studies of the citrate-responsive sensor histidine kinase, and took advantage of the same conformational changes proposed to propagate from the binding domain to the catalytic domain. Following extensive engineering based on a combination of structure guided mutagenesis and directed evolution, we produced an inverse-response biosensor (ΔF/F_min_ ~ 18) designated Citroff1 and a direct-response biosensor (ΔF/F_min_ ~ 9) designated Citron1. We report the x-ray crystal structure of Citron1 and demonstrate the utility of both biosensors for qualitative and quantitative imaging of steady-state and pharmacologically-perturbed citrate concentrations in live cells.

## Introduction

Citrate is primarily known as the first major intermediate in the tricarboxylic acid (TCA) cycle, but in recent years there has been a growing appreciation of the wider range of roles for citrate in biological processes including inflammation, cancer, and insulin secretion, among others^1^. Citrate is produced in the mitochondria by the enzyme citrate synthase (CS), and is metabolized into isocitrate by the enzyme aconitase 2 (ACO2) in the next step of the TCA cycle. Citrate can be transported from the mitochondria to the cytosol by the citrate carrier (CiC; SLC25A1), or transported into the cell from the bloodstream by a plasma membrane-specific variant (pmCiC; SLC13A5) of the mitochondrial CiC. Once in the cytosol, citrate can be cleaved by ATP citrate lyase (ACLY) into acetyl-CoA and oxaloacetate. Acetyl-CoA is the substrate for various biochemical processes including fatty acid synthesis and histone acetylation for the epigenetic modifications of gene expression^2^, and oxaloacetate plays a key role in various biochemical processes including amino acid biosynthesis. As some of citrate’s many roles are associated with the mitochondrial pool, and others are associated with the cytoplasmic pool, methods that can enable the accurate determination of the concentration in both subcellular compartments are of high importance.

Much remains to be learned about the roles of citrate, as published reports support apparently contradictory roles for this key metabolite in cancer cells. Huang et al. have noted that citrate can have both anti-tumor and tumor-promoting effects, depending on its local concentration^2^. For example, despite the decreased reliance on the TCA cycle in tumor cells (the Warburg effect)^3^, some cell types appear to have an increased cytosolic pool of citrate that may serve as a source of acetyl-CoA to be used for increased fatty acid biosynthesis^4^. Conversely, other evidence suggests that a decrease in cytosolic citrate is associated with tumor aggressiveness, possibly due to increased resistance to apoptosis as a consequence of decreased acetyl-CoA dependent protein acetylation^5^. As these examples demonstrate, citrate is central to the dysregulated metabolism of cancer cells, and enzymes that produce (i.e., CS), transport (i.e., CiC), and metabolize (i.e., ACLY and ACO2), citrate have been proposed as potential targets for new therapeutic approaches to cancer^2,4^. Further investigation of the role of citrate in tumor development will require new methods that could be used for sensitive detection of citrate in subcellular compartments of cells in *in vivo* tumor models.

In addition to its roles in cancer cells, citrate functions as a regulatory metabolite in endocrine cells such as beta cells^6^ and immune cells such as dendritic cells and macrophages^7^. Previous studies suggest that citrate (and isocitrate) exported by CiC to the cytosol can function as a signalling molecule to regulate downstream insulin secretion in beta cells^8,9^. Furthermore, activated dendritic cells and macrophages undergo a metabolic reprogramming that leads to the accumulation of citrate, which, in turn, regulates the immune cell functions through carbohydrate and fatty acid metabolism and citrate-derived acetylation of histones^10^. Techniques that enable spatially and temporally resolved imaging of citrate concentrations would facilitate the understanding of its central functions in these metabolism-based signaling processes.

A variety of synthetic fluorescent indicators for citrate have been reported, including ones based on indicator displacement assays^11–13^, turn-on fluorescence complexation^14^, competitive binding of metal ions such as Eu^3+^ and Pb^2+^ (Refs.^15,16^), and aggregation induced emission^17^. These indicators have proven useful for the quantitative analysis of citrate in urine but none of these designs are practical for measurement of citrate in the mitochondria and cytosol of living cells. Challenges with their use in such applications include delivery into cells, competition from other endogenous small molecules, potential toxicity, and the lack of control over subcellular location.

For *in vitro* or *in vivo* application in tissue, an attractive alternative to synthetic indicators are fully genetically encoded proteinaceous biosensors^18^. Genetically encoded biosensors are engineered proteins composed of a fusion between a fluorescent protein (FP), such as *Aequorea victoria* green FP (GFP), and a sensing domain that undergoes a conformational change upon binding its cognate ligand of interest, or in response to some other biological parameter of interest. This conformational change in the sensing domain is propagated to the FP fluorophore environment, leading to a change in fluorescence intensity. First demonstrated in 1999 by Tsien and coworkers for the construction of Ca^2+^ and Zn^2+^ biosensors^19^, this general approach has now been expanded to a multitude of other ligands of interest^18^. Advantages of such biosensors include their ability to be delivered relatively non-invasively in the form of their DNA genes, high specificity conferred by the molecular recognition properties of the sensing domain, low toxicity due to being protein based, and the ability to be genetically targeted to subcellular compartments.

A key challenge in developing a genetically encoded biosensor is first identifying a protein that both specifically binds to a target ligand of interest and undergoes substantial conformational change upon binding. A protein that satisfies both of these criteria are sensor histidine kinases (SHK), which are one of the two proteins in a bacterial two-component regulatory system (with the other protein being the response regulator)^20^. A typical SHK is a homodimer protein with an N-terminal periplasmic sensory domain that has both its N-terminus (TM1) and C-terminus (TM2) linked to transmembrane α-helices (**Fig. 1a**). Linked to TM2 are several intracellular domains, including the C-terminal histidine kinase (HK) catalytic domain^21^. Necessarily, binding of a cognate ligand to the periplasmic sensory domain leads to a conformational change that propagates across the membrane and modulates the activity of the C-terminal HK domain^21–23^. Various models have been proposed for these membrane-spanning structural changes including the “piston” model, in which the two helices are vertically displaced relative to each other, the “scissor blade” model, in which the helices undergo a diagonal scissor-like displacement, and the “rotating helix” model in which one helix is proposed to rotate relative to the other^21^. Regardless of the precise mechanism, it is evident that ligand binding to the periplasmic domain must lead to substantial conformational changes in the transmembrane portion of the HKS.

**Figure 1.**
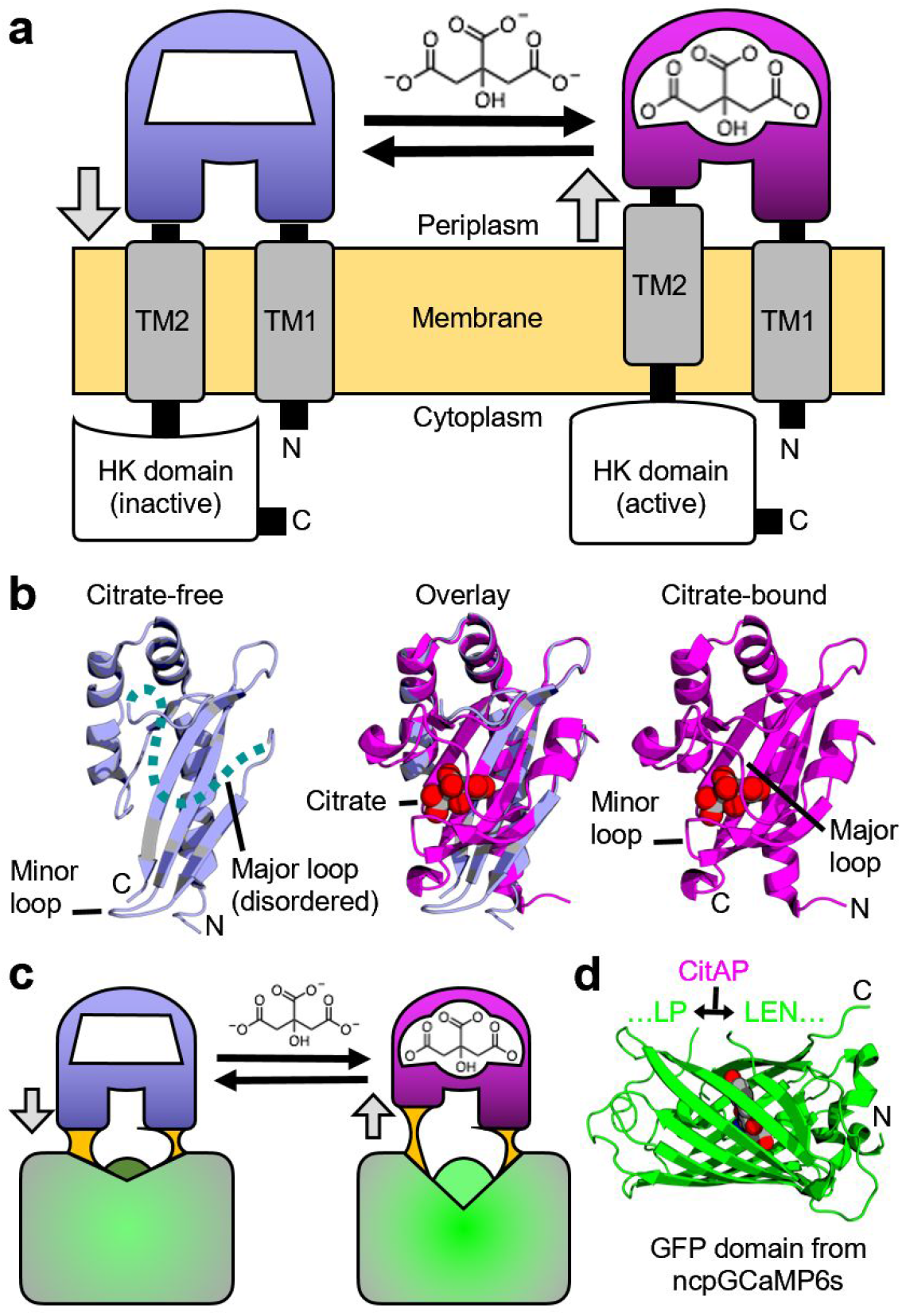
Rationale for the design of a single FP-based citrate biosensor. **a** Schematic representation of *Klebsiella pneumoniae* SHK CitA, which is composed of a periplasmic citrate-binding domain (CitAP; light blue, unbound; magenta, bound), connected to transmembrane helices at both its N- (transmembrane helix 1, TM1) and C-termini (transmembrane helix 2, TM2). TM2 is, in turn, connected to an intracellular HK catalytic domain^26^. **b** The structures of citrate-free CitAP (left; light blue; PDB ID 2V9A)^25^, citrate-bound CitAP (right; magenta; PDB ID 2J80)^25^, and a superposition of the citrate-free and -bound structures (middle). **c** We hypothesized that the piston-type conformational motion at the CitAP termini could be communicated to GFP to allosterically control the chromophore environment and its fluorescent brightness. In this way, the CitAP domain could serve as the basis of construction of a genetically encoded citrate biosensor. **d** To realize this biosensor design, we inserted CitAP into GFP by replacing the CaM-RS20 domain of ncpGCaMP6s^27^ with CitAP.

One relatively well characterized SHK is the citrate-responsive CitA protein from *Klebsiella pneumoniae^22,24,25^*. X-ray crystal structures are available for the Per-Arnt-Sim (PAS)-fold citrate-binding periplasmic domain (CitAP) both with (PDB IDs 1P0Z^24^ and 2J80)^25^ and without (PDB ID 2V9A) ^25^ citrate (**Fig. 1b**). As described by Sevanna et al.^25^, the major loop of CitAP is disordered in the citrate-free structure, but ‘wraps over’ the citrate binding site in the bound state. In addition, there is substantial movement of a minor loop that ‘folds in’ on the citrate binding site in the bound state. This ‘folding in’ of the minor loop is associated with a compaction of the central β-sheet that pulls the C-terminus away from the transmembrane domain and causes a piston-like “pull” on TM2. This model is also supported by NMR studies of a truncated and membrane embedded form of CitA^22^.

Honda and Kirimura previously reported a genetically encoded biosensor, designated CF98, based on insertion of circularly permuted (cp) GFP into the CitAP domain of CitA^28^. To develop CF98, Honda and Kirimura made a series of nine prototype constructs in which circularly permuted cpGFP was systematically inserted at positions within the minor loop of CitAP (**Fig. 1b** and **Fig. S1ab**. The most promising construct (CF98), was based on insertion of cpGFP between residues 98 and 99 of CitAP (all numbering as in PDB ID 2J80) ^25^, and exhibited an intensiometric fluorescence increase (ΔF/F, calculated as (F_max_ − F_min_)/ F_min_) of approximately 0.7. CF98 was used to detect intracellular citrate concentration changes in a suspension of *E. coli* treated with citrate. To the best of our knowledge, there have been no reports of using this biosensor to visualize citrate concentrations in eukaryotic cells. In other work, Ewald et al. reported a series of genetically encoded Förster resonance energy transfer (FRET)-based biosensors, composed of a cyan FP and yellow FP fused to the termini of CitAP, that exhibit up to a 55% increase in emission ratio upon binding citrate ^29^. These biosensors were originally used to monitor citrate in *E. coli* after starvation and were later used to monitor citrate oscillations in glucose-treated islet β-cells^30^.

With the growing recognition of the central importance of citrate in healthy and diseased cells^1^, and the need for new tools to probe the concentration of citrate in tissues, we have developed a new generation of high performance intensiometric direct-response (Citron) and inverse-response (Citroff) genetically encoded biosensors for citrate. Further motivation came from the growing recognition that, with appropriate optimization and directed evolution, high performance single-FP based biosensors can be engineered with ΔF/F > 10 (Refs.^31,32^), or even > 100 (Refs.^33,34^). Such large fluorescence changes, combined with high fluorescent brightness and ligand affinity tuned to the relevant physiological concentration, are critical if such biosensors are to be robust and effective tools for tissue-based imaging experiments.

## Results

### Development of citrate biosensors Citron1 and Citroff1

To construct a genetically encoded biosensor for citrate, we replaced the calmodulin (CaM)-RS20 domain of ncpGCaMP6s^27^ (from pcDNA-ncpGCaMP6s, Addgene plasmid #113674) with residues 4 - 133 of the CitAP domain of *Klebsiella pneumoniae* CitA (**Fig. 1c,d** and **Fig. S1cd**). From this point on, residues are numbered according to the full length biosensor constructs, as shown in **Fig. S2**. The initial variant with linker sequences copied from ncpGCaMP6s (linker 1 (L1) = L145-P146 and linker 2 (L2) = L277-E278-N279) was non-fluorescent. To rescue the fluorescence, we randomized each of the two linkers individually and performed colony-based screening. Screening of a library (~1000 colonies) in which both residues of L1 were randomized, led to the identification of ~50 fluorescent colonies with relatively high brightness. These colonies were picked and cultured overnight at 37 C in 4 mL LB media. The bacteria were lysed, the protein extract dispensed into wells of a 384-well plate, and a microplate reader was used to evaluate fluorescence changes upon citrate binding. This screening led to a variant, designated Citron0.1 (L1 = L145M-P146V), with a ΔF/F, calculated as (F_max_ - F_min_)/ F_min_, ~ 0.2 direct-response (that is, an increase in fluorescence upon citrate binding), and a second variant, designated Citroff0.1 (L1 = L145W-P146Q), with ΔF/F ~ 1.9 inverse-response (that is, a decrease in fluorescence upon citrate binding). A similar randomization and screening of L2 was then performed for both variants, leading to the identification of Citron0.2 (ΔF/F ~ 0.6; L2 = L277S-E278N-N279L), and Citroff0.2 (ΔF/F ~ 4.5; L2 = L277P-E278N-N279F).

Following linker optimization, the two variants were further optimized in parallel by directed evolution, which involved screening of randomly mutated libraries generated by error-prone polymerase chain reaction of the entire biosensor gene. After each round, the most promising variants were used as the template for the next round of library construction and screening. Following 10 rounds of directed evolution, the best direct-response biosensor (with 13 mutations in GFP and 8 mutations in CitAP; **Fig. S2**) had ΔF/F ~ 9 *in vitro* (**Fig. 2a** and **Table S1**) and was named as Citron1. For the inverse-response biosensor, three rounds of directed evolution led to a variant (with 3 mutations in GFP and 1 mutation in CitAP; **Fig. S2**) that exhibited ΔF/F ~ 18 *in vitro* (**Fig. 2b** and **Table S1**) and was designated Citroff1. Further screening did not produce any variants with improved performance. The dynamic range of both biosensors are substantially higher than the ΔF/F ~ 1.1 that we measured for purified CF98 under identical experimental conditions (**Fig. S3a**). The molecular brightness of the bright states of Citron1 (+ citrate) and Citroff1 (− citrate), calculated as the product of extinction coefficient and quantum yield, are 21 and 25, respectively (**Table S1**). For reference, EGFP has a brightness of 37.5 (Ref.^35^) and the Ca^2+^-bound state of GCaMP6s has a brightness of 41.8 (Ref.^36^).

**Figure 2.**
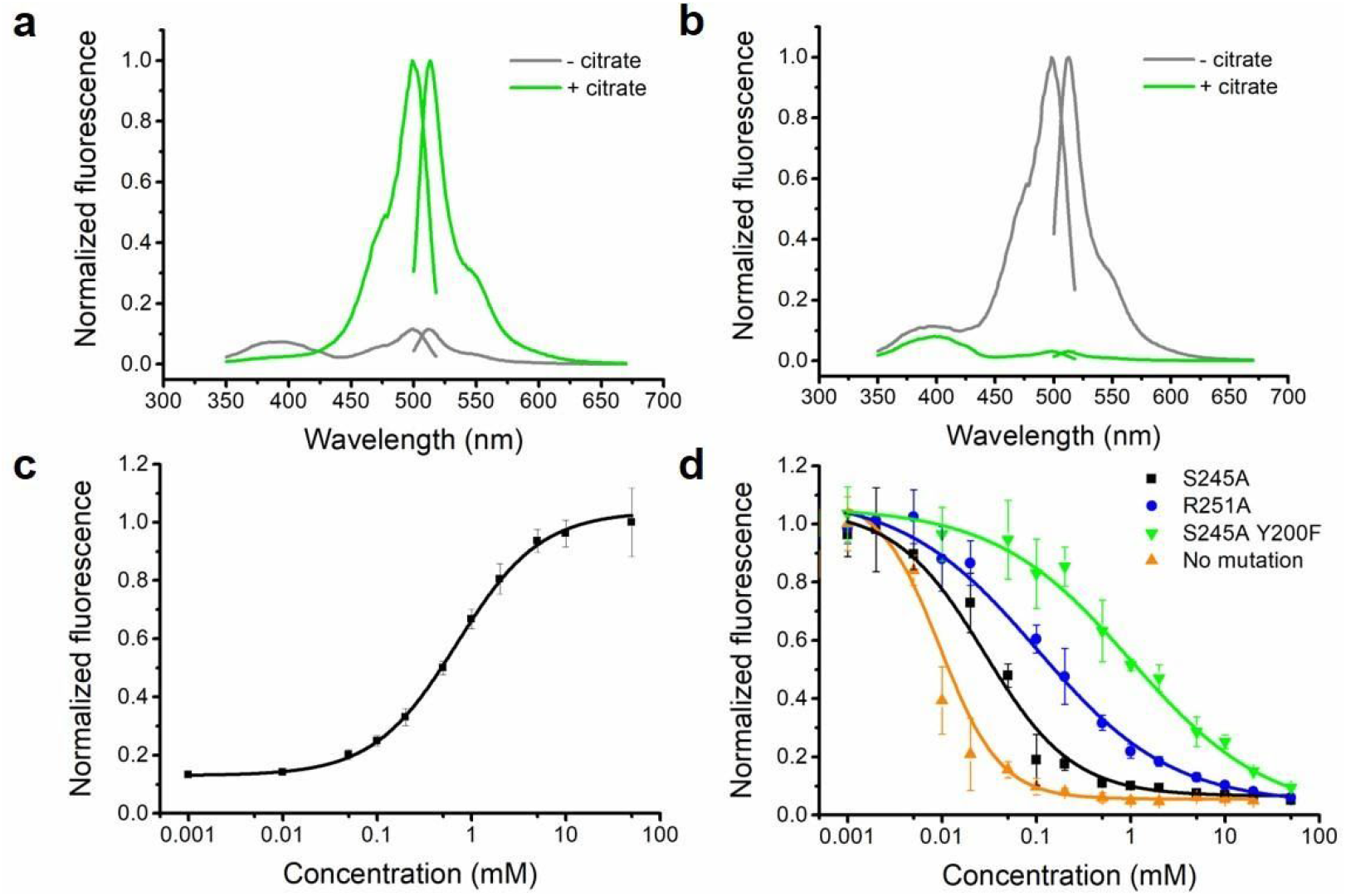
*In vitro* characterization of new citrate biosensors. **a,b** Normalized excitation and emission spectra of purified Citron1 (**a**) and Citroff1 (**b**) in the presence and absence of 20 mM citrate. **c,d** *In vitro* citrate titration curves of purified Citron1 (**c**) and Citroff1 variants (**d**). Error bars represent standard deviation of triplicates.

### *In vitro* characterization and affinity tuning

Determination of the citrate binding affinities of the two biosensors revealed that the *K*_d_ of the direct-response biosensor, Citron1, is 1.1 mM (**Fig. 2c**), which is a substantially lower affinity than that of CitAP itself (*K*_d_ = 5.5 μM at pH 7)^37^. This decreased affinity is attributed to mutations in the CitAP domain that were accumulated during directed evolution (**Fig. S2**). Specifically, mutation E223V is immediately adjacent to the citrate binding pocket (equivalent to E80V as numbered in PDB ID 2J80)^25^ and is likely to directly affect citrate binding (**Fig. S4**). However, as the intracellular citrate concentration in mammalian cells ranges from hundreds of micromolar to several millimolar, Citron1 should be particularly responsive to physiologically relevant concentration changes. We determined that CF98 demonstrates an even lower binding of affinity of ~ 9 mM which is likely to make it poorly responsive to physiologically relevant changes in citrate concentration (**Fig. S3b**).

In contrast to Citron1, Citroff1 retains the high binding affinity of CitAP (**Fig. 2d**). In an effort to fine tune the affinity of Citroff1 into a physiologically relevant concentration range, we used site-directed mutagenesis to introduce mutations (Y199F, T201A, S227A, S244A, S248A, and S267A) that were expected to directly or indirectly disrupt interactions between the protein and citrate, based on the crystal structure of CitAP (PDB ID 2J80)^25^. Mutations that have previously been reported to lower the binding affinity (R192A, K220A, K235A and R250A) were also assessed^38^. Single mutations S244A and R250A resulted in biosensors with substantially lower affinity (*K*_d_ = 35 μM and 153 μM, respectively) while maintaining a large dynamic range (**Fig. 2d** and **Table S2**). A combination of S244A and Y199F further increased the *K*_d_ to 965 μM but with a slightly lowered ΔF/F of ~ 11 (**Fig. 2d** and **Table S2**). Other single or pairwise combinations of mutations did not yield variants that had both appropriate affinities and large fluorescence responses (**Fig. S5**).

### pH dependence and non-binding control biosensors

To determine the pH dependence of the response of Citron1 and Citroff1 the fluorescence was measured in the presence and absence of citrate at various pHs (**Fig. S6a,b**). Both biosensors exhibit reasonably good pH stability over the physiologically relevant pH range. In terms of specificity, the two citrate biosensors showed negligible response to several potentially interfering metabolites at high concentration (20 mM) (**Fig. S6c,d**). This excellent specificity is similar to that of the previously reported citrate biosensors that employed CitAP as the sensing domain^28,29^. In a previous study, a bound sodium ion (Na^+^) was observed in the crystal structure of CitAP, which might potentially affect the citrate-induced conformational change.^24^ The dependence of Citron1 and Citroff1 on Na^+^ was examined and no obvious Na^+^-dependent fluorescence change was found (**Fig. S7**).

While both Citron1 and Citroff1 are only modestly pH sensitive through the physiologically relevant pH range, the possibility of artifactual fluorescence changes due to changes in pH is a persistent concern when performing cell imaging with genetically encoded biosensors. Accordingly, we created citrate-insensitive (but pH-sensitive) control constructs by introducing two key mutations (R209A and H212A, equivalent to CitAP R66A and H69A, respectively; **Fig. S2** and **Fig. S8a**) to disable citrate binding. These variants, named as CitronRH and CitroffRH, showed no response to citrate (**Fig. S8b,c**) but pH dependence that was similar to Citron1 and Citroff1 in the citrate-free states (**Fig. S8d,e**). These variants are appropriate controls for evaluating the possible contribution of pH changes to fluorescence responses observed during cell imaging experiments.

### Crystal structure of Citron1

In an effort to obtain molecular insight into the structure and mechanism of these citrate biosensors, we crystallized Citron1 in the citrate-bound state. The Citron1 crystal structure was determined to 2.99 Å resolution using molecular replacement (**Fig. 3, Table S3**). The overall structure is represented in **Fig. 3a**, with all mutations labelled. Within the biosensor structure, the CitAP domain and its citrate binding pocket are essentially identical to previously determined CitAP structures^24,25^.

**Figure 3.**
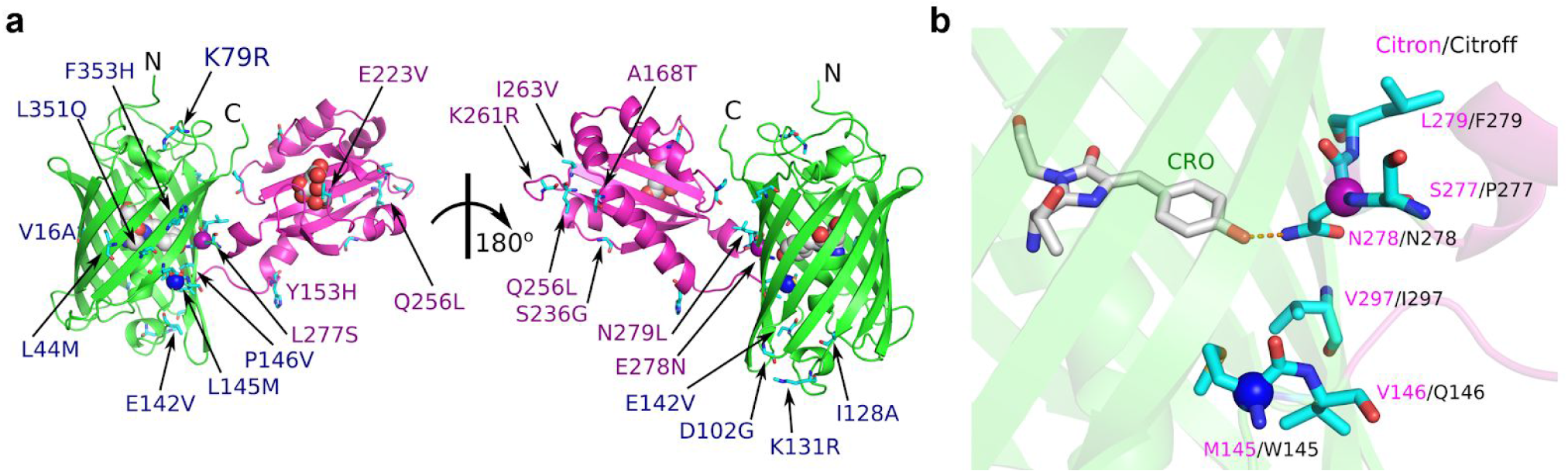
Crystal structure of Citron1. **a** Overall representation of the Citron1 structure with the position of all mutations indicated. The CitAP domain with citrate is coloured in magenta, the cpGFP domain is coloured in green. The chromophore, citrate, and the Cɑ of Met145 (blue) and Asn278 (magenta) are represented as spheres. **b** Zoom-in view of the hydrogen bonding interaction between Asn278 and the chromophore. Additional residues in the vicinity of Asn278 of Citron1 are shown and labeled with magenta text, with the identity of the corresponding residue of Citroff1 labelled with black text.

Inspection of the molecular interactions between the GFP chromophore and its immediate environment reveals a possible mechanism for the citrate-dependent fluorescence modulation. Specifically, we observe a hydrogen bond between the GFP chromophore phenolate group and the side chain of Asn278 in linker 2 (**Fig. 3b**). This interaction is likely to stabilize the fluorescent phenolate form (deprotonated form), resulting in bright fluorescence in the citrate-bound state. Presumably, in the citrate-free state, Asn278 is positioned in a different conformation such that this hydrogen bond interaction is not present, and the dimly fluorescent phenol form (protonated form) of the chromophore predominates. A very similar mechanism has been previously suggested for the red Ca^2+^ biosensor K-GECO1 (Ref. 39). In the Ca^2+^-bound state of K-GECO1, Asn32 is positioned similarly to Asn278 of Citron1, and is similarly engaged in a hydrogen bond with the chromophore. In the case of Citroff1, we speculate that Asn278 stabilizes the phenolate form of the chromophore through a hydrogen bond interaction in the citrate-free state, but not in the citrate-bound state.

### Quantification of citrate concentrations in HeLa cells

To evaluate the utility of Citron1 and Citroff1 for visualization of intracellular citrate concentration changes, we expressed each construct in HeLa cells (ATCC CCL-2). The genes were expressed under a CMV promoter with either no targeting (that is, cytoplasmic localization), or as a fusion to two copies of a mitochondrial-targeting sequence^40^ (that is, mitochondrial localization). As shown in **Fig. 4a** the biosensors exhibited strong fluorescence intensity in both the cytoplasm and mitochondria of HeLa cells. Artificial changes in the intracellular citrate concentration were achieved by permeabilization of the cell membrane with digitonin followed by citrate addition to the imaging buffer. Substantial fluorescence changes were observed for both Citron1 and Citroff1, and the changes were similar for both the cytoplasmic and mitochondria-localized biosensors (**Fig. 4b** and **Fig. S9a**). No substantial fluorescence changes were observed when cells expressing the control variants, CitronRH and CitroffRH, were subjected to the same treatments (**Fig. S10a,b**). These results demonstrate that Citron1 and Citroff1 remain responsive to citrate concentration changes when expressed in live cells.

**Figure 4.**
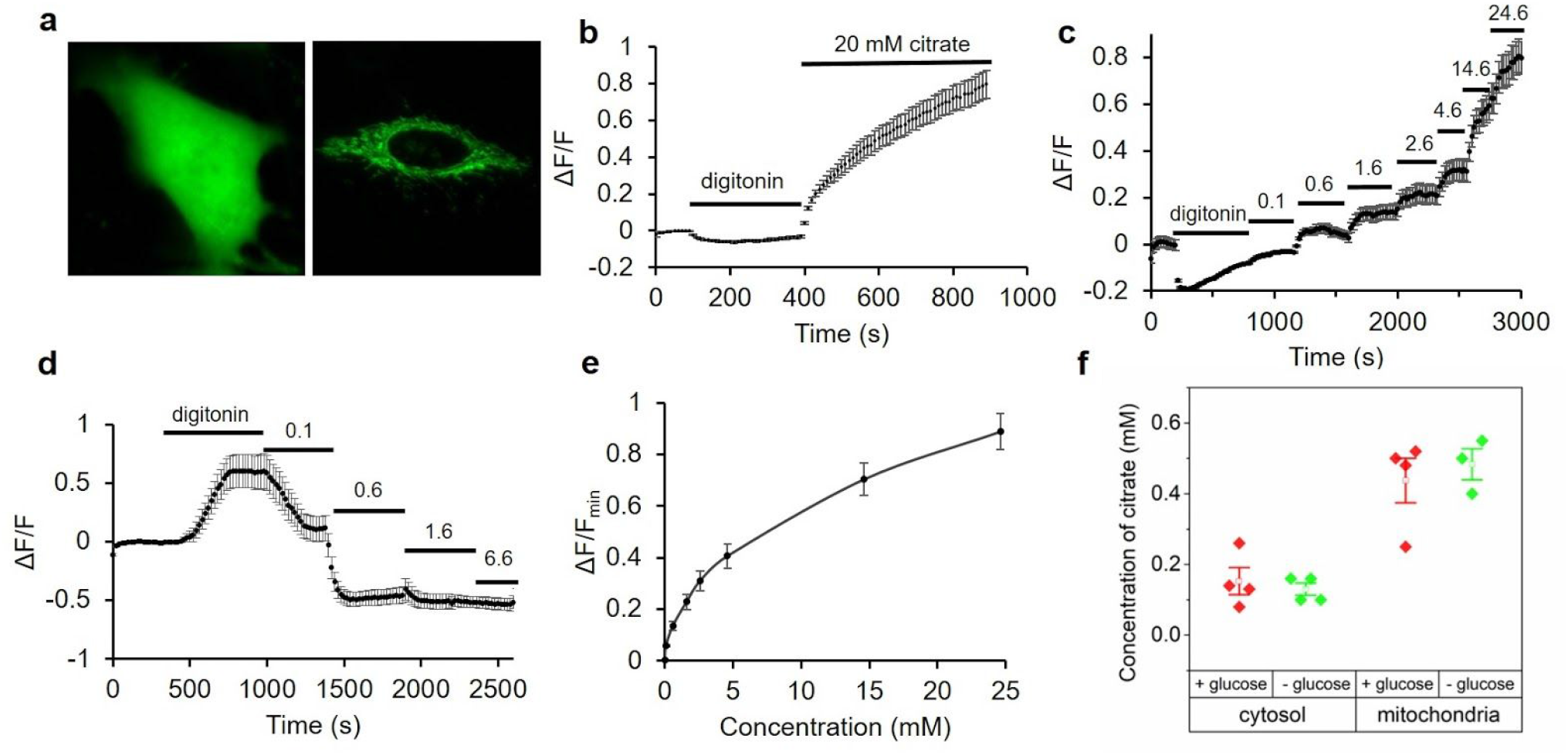
Expression of citrate biosensors in HeLa cell and permeabilization tests. **a** Representative fluorescence images of HeLa cells expressing Citron1 in the cytosol (left panel) and mitochondria (right panel). **b** Fluorescence intensity of Citron1 in the cytosol in response to treatment with digitonin and citrate (n = 78 cells). The analogous chart for mitochondrial Citron1 is provided as **Fig. S9a**. **c, d** *In situ* titration of Citron1 (**c**) and Citroff1 (**d**) in the cytosol. Digitonin was added as indicated and the concentration of citrate in the external buffer is indicated in millimolar (n = 128 for Citron1 and n = 26 for Citroff1). **e**. *In situ* titration curve of Citron1 in the cytosol averaged from 128 cells in panel **c**. The analogous chart for Citroff1 is provided as **Fig. S9b**. Error bars represent s.e.m for panels **b**-**e**. **f** Quantification of citrate concentration in the cytosol and mitochondria with or without 5.5 mM glucose in the buffer. Each dot is quantified using the average signal from tens of cells in a single experiment. Center box and whiskers represent the average and s.e.m. of the four data points, respectively.

To perform an *in situ* titration to quantify the citrate concentration in the cytosol, cell membranes were permeabilized with digitonin and various concentrations (0.1 to ~ 20 mM) of citrate were added to the imaging buffer (**Fig. 4c,d**). The average response from cells expressing Citron1 or Citroff1 vs. concentration of citrate was plotted to provide an *in situ* calibration curve (**Fig. 4e** and **Fig. S9b**). To quantify the concentration of citrate in cells, we acquired a fluorescence image of individual intact cells, then acquired a second image following treatment with digitonin to fully deplete intracellular citrate, and then acquired a third image after adding a saturating concentration of citrate (20 mM). The initial fluorescence intensity of intact cells (first image) was normalized according to the intensities at 0 mM (second image) and 20 mM citrate (third image) and the initial concentration determined using the previously acquired calibration curve (**Fig. 4e** and **Fig. S9b)**. Using this approach, the average citrate concentration in the cytosol and mitochondria of HeLa cells conditioned in HEPES buffered HBSS solution with 5.5 mM glucose were determined to be 0.15 ± 0.07 mM and 0.44 ± 0.13 mM, respectively, using Citron1 (four replicates each). In the absence of glucose in the same buffer, citrate concentration remained essentially constant both in the cytosol and mitochondria (**Fig. 4f**). Quantification using Citroff1 gave concentrations (0.12 ± 0.01 mM in the cytosol and 0.45 ± 0.19 mM in the mitochondria) consistent with the Citron1 measurements. The experimentally determined cytosolic citrate concentration is comparable to that previously reported for Hep G2 cells (168.0 ± 51.0 μM)^41^ and human blood plasma (100 - 150 μM)^42^.

### Physiological and pharmacological alteration of subcellular citrate concentration

Next, we explored the use of Citron1, Citroff1 and the affinity variants for imaging of changes in citrate concentration induced by changes in culture media composition or by pharmacological treatments. Both Citron1 and Citroff1 biosensors demonstrated larger *K*_d_ in cultured cells than that *in vitro* (5.7 mM *in situ* (**Fig. 4e**) vs. 1.1 mM *in vitro* for Citron1 and 130 μM *in situ* (**Fig. S9b**) vs. 5 μM *in vitro* for Citroff1). Among the inverse-response biosensors, Citroff1 has optimal *in situ K*_d_ for imaging citrate dynamics in the cytosol where citrate concentration is determined to be 0.12-0.15 mM. Citroff1 S245A, with slightly lower affinity than Citroff1, is presumably suitable for imaging citrate in the mitochondria where citrate concentration was ~0.45 mM. To investigate glucose-induced changes in intracellular citrate concentration, Citron1- and Citroff1-expressing HeLa cells were kept in a glucose-free buffer for 0.5 - 1 h. Upon glucose addition, both Citron1 and Citroff1, targeted to either the cytosol or mitochondria, exhibited fluorescence intensity changes consistent with an increased citrate concentration (that is, increased fluorescence intensity for Citron1 (**Fig. 5a**) and decreased signal for Citroff1 (**Fig. S11b**)). The cytosolic citrate concentration substantially increased up on glucose treatment as revealed using both Citron1 and Citroff1. For mitochondrial citrate, treatment with glucose induced an initial transient decrease followed by a slower increase as reported by the mitochondria-localized Citron1. Consistent results were obtained with mitochondrially targeted Citroff1 S245A (with *K*_d_ = 35 μM vs. *K*_d_ = 5 μM for Citroff1), though the initial transient decrease was not as pronounced (**Fig. S11d**). Similar experiments using the control variants, CitronRH and CitroffRH, confirmed that the observed signal changes were attributable to changes in citrate concentration rather than other potentially confounding factors such as pH changes (**Fig. S11**).

**Figure 5.**
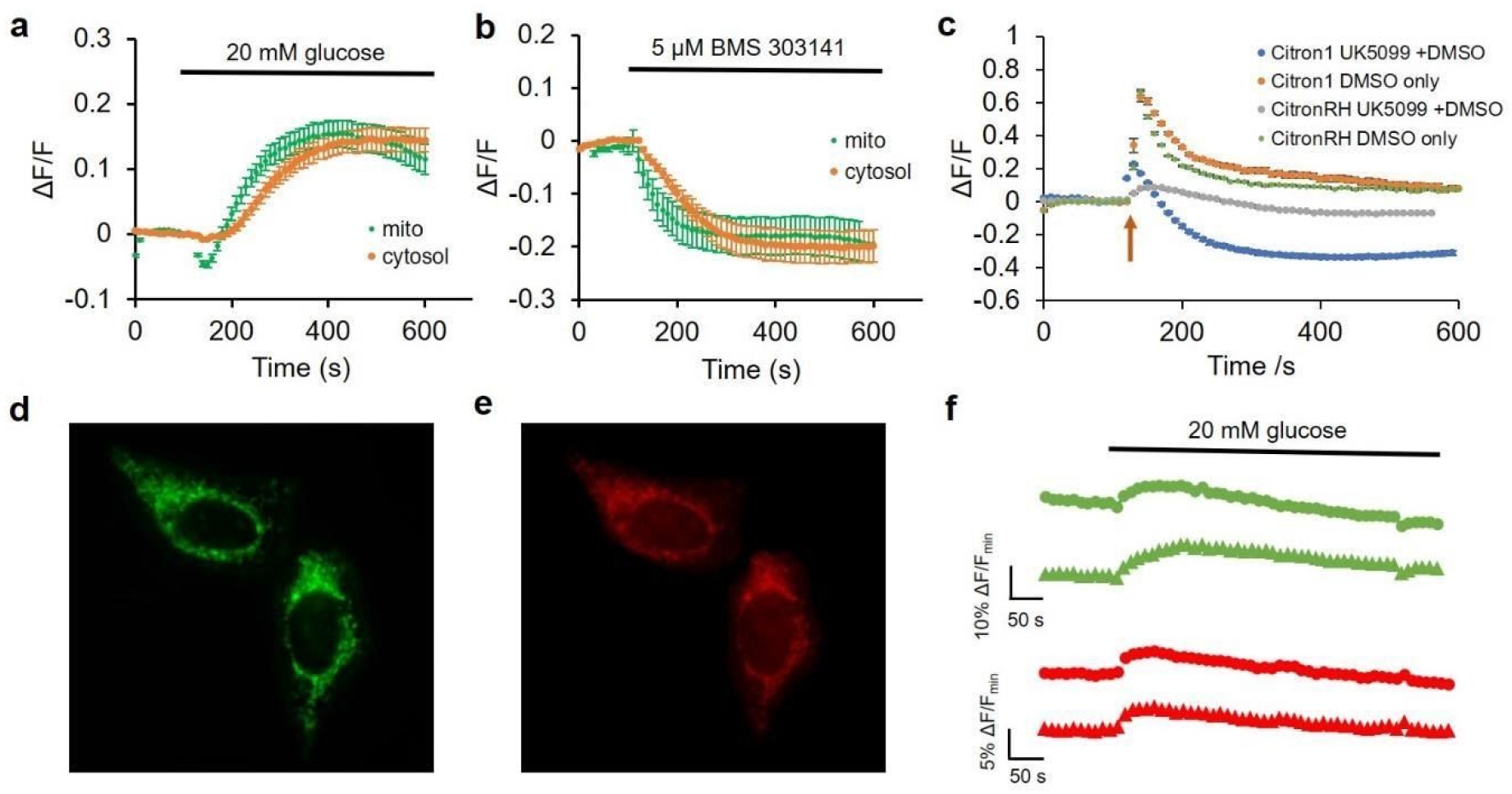
Use of Citron1 for imaging of citrate concentration changes induced by changes in glucose or pharmacologically in HeLa cells. **a** Glucose-induced citrate concentration changes detected with cytosolic (orange trace, n = 32) and mitochondrial (green trace, n = 28) Citron1. **b** BMS 303141-induced citrate concentration changes detected by cytosolic (n = 22) and mitochondrial (n = 24) Citron1. **c** UK-5099 (in DMSO)-induced citrate concentration changes detected by Citron1 (blue trace, n = 36) expressed in mitochondria. Control experiments include Citron1 + DMSO (orange trace, n = 23), CitronRH + UK-5099 (grey trace, n = 38), and CitronRH + DMSO (green trace, n = 23). The arrow indicates the addition of UK-5099 or DMSO solutions. Error bars in a-c represent s.e.m.. **d-f** Dual color imaging of citrate and ATP concentration changes in mitochondria using Citron1 and MalionR^43^. **d,e** Representative fluorescence images of cells co-expressing Citron1 (**d**) and MalionR (**e**). **f** Representative glucose-induced citrate and ATP concentration changes reported by Citron1 (green traces) and MalionR (red traces).

The observed increases in cytoplasmic and mitochondrial citrate concentrations following glucose treatment are consistent with increased glucose availability fueling the TCA cycle, resulting in an increase in TCA intermediates including citrate. Notably, citrate concentration changes occurred approximately 50 s earlier in the mitochondria compared to the cytosol (**Fig. 5a**, consistent with the expected production of citrate by citrate synthase in mitochondria^10^.

We next used Citron1 and Citroff1 to investigate changes in citrate concentration associated with inhibition of ACLY by the inhibitor BMS-303141 (Ref. ^44^). Using cytosolic Citron1 and Citroff1, we observed a transient slight increase of citrate concentration, followed by a sustained decrease in concentration(**Fig. S12a, b**). The initial transient increase in mitochondrial citrate concentration was most pronounced when using Citroff1 S245A (**Fig. S12d**), but less apparent with Citron1 (**Fig. 5b, Fig S12c**). These differences in biosensor response are most likely due to differences in *K*_d_ for Citron1 vs. Citroff1 vs. Citroff1 S245A (1100 μM, 5 μM, and 35 μM, respectively). Mechanistically, the transient citrate accumulation is to be expected upon ACLY inhibition, due to cessation of citrate to acetyl-CoA conversion. The subsequent sustained decrease in mitochondrial citrate concentration may be due to the suspended production of oxaloacetate catalyzed by ACLY and downstream malate catalyzed by malate dehydrogenase^45^ in the cytosol. Since the mitochondrial CiC imports malate to the mitochondria in exchange for citrate export^46,47^, the reduced concentration of cytosolic malate could potentially suppress citrate export and therefore cause a decrease in the concentration of citrate in the cytosol. Parallel experiments with the control variants CitronRH and CitroffRH confirmed that the observed changes in fluorescence were induced by citrate concentration changes (**Fig. S12**). Taken together, our results demonstrate that imaging of citrate concentration dynamics with Citron1 and Citroff1 provides a sensitive method for the detection of ACLY activity that could potentially be used in cell-based screens for ACLY inhibitors^4^.

To investigate changes in citrate concentration associated with pharmacological manipulation of its upstream metabolic regulator, mitochondrial pyruvate carrier (MPC), we applied MPC inhibitor UK-5099 to HeLa cells expressing the mitonchondria-localized citrate biosensors^48^. We hypothesized that inhibition of MPC would block pyruvate uptake into mitochondria and lead to decreased flux of the TCA cycle and a lower citrate concentration. As shown in **Fig. 5c** (blue trace), treatment with UK-5099 (1 mM stock in 10% DMSO; diluted 1/40 to give a final concentration of 25 μM) resulted in a transient 20% fluorescence increase of Citron1, followed by a sustained decrease. In parallel control experiments, we found that DMSO alone elicited even larger transient fluorescence increases for both Citron1 and CitronRH in mitochondria (**Fig. 5c**, green trace), presumably due to a pH increase. When cells expressing CitronRH were treated with both DMSO and UK-5099, the large transient fluorescence change was suppressed (**Fig. 5c**, grey trace). These results suggested the observed changes in Citron1 fluorescence upon treatment of cells with DMSO and UK-5099 are primarily due to changes in citrate concentration, with a smaller contribution from pH changes. These results are consistent with UK-5099-dependent inhibition of pyruvate uptake into mitochondria and suggest that Citron1 could be potentially used for indirect monitoring of pyruvate uptake and TCA cycle activity in mitochondria. This example also illustrates the importance of non-binding control constructs to decouple true from artifactual fluorescence changes.

### Concurrent imaging of Citrate and ATP dynamics in HeLa Cells

To explore the possibility of using the new citrate biosensors for multicolor and multiparameter metabolite imaging, we co-expressed green fluorescent Citron1 (**Fig. 5d**) with the red fluorescent ATP biosensor, MalionR (**Fig. 5e**)^43^, both targeted to the mitochondria. Upon treatment of starved (glucose-free) HeLa cells with 20 mM glucose, we observed a change in MalionR fluorescence that was consistent with an increase in ATP concentration, as has been previously reported^49^. Similar to our previous results (**Fig. S11c**), Citron1 again reported an initial small dip and then a large increase in mitochondrial citrate (**Fig. 5f**). Overall, these results support the notion of tight coupling between ATP and citrate concentrations in mitochondria^50^.

### Imaging citrate in INS-1 beta cells

Citrate export from the mitochondria plays a critical role in regulation of insulin secretion from pancreatic beta cells^8^. To investigate the utility of the citrate biosensors in beta cells, we expressed Citron1 in INS-1 rat insulinoma cells (**Fig. 6a**) and acquired *in situ* calibration curves (**Fig. 6b,c**). Using the *in situ* calibration curves, we determined the concentration of citrate in the cytosol and mitochondria of INS-1 cells, conditioned in Krebs-Ringer buffer, to be 2.93 ± 0.09 mM and 2.81 ± 0.24 mM, respectively (**Fig. 6d**). These concentration values are substantially higher than those of HeLa cells (0.15 ± 0.07 mM and 0.44 ± 0.13 mM). The higher concentrations of citrate are consistent with a higher flux through the TCA cycle, as further supported by the higher oxygen consumption rates (OCRs) of INS-1 cells compared to that of HeLa cells ^49^. Addition of 20 mM glucose increased both cytosolic and mitochondrial citrate concentration to ~25 mM (**Fig. 6d-f**). This result is in agreement with a previous study in which it was demonstrated that stimulatory glucose can result in an approximately 8-fold increase in the concentration of citrate^51^.

**Figure 6.**
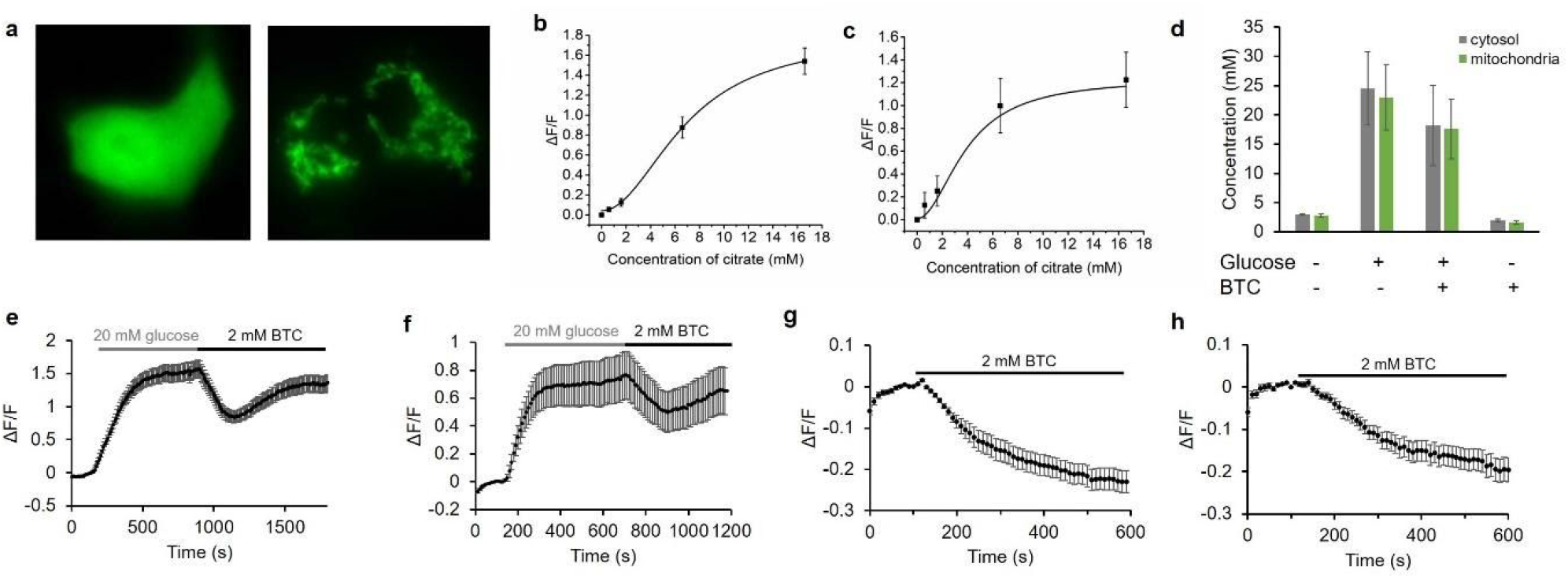
Characterization of Citron1 in INS-1 cells. **a** Representative fluorescence images of INS-1 cells expressing Citron1 in the cytosol (left panel) and mitochondria (right panel). **b,c** *In situ* titration curve of Citron1 in the cytosol (**b**, n = 62) and mitochondria(**c**, n = 34). **d** Citrate concentration in the cytosol (grey) and mitochondria (green) in Krebs-Ringer buffer with or without 20 mM glucose/2 mM BTC treatment. Each quantification result is averaged from triplicates. **e,f** Glucose and BTC induced citrate changes in the cytosol (**e**, n = 49) and mitochondria (**f**, n = 14). **g,h** BTC induced citrate changes in the cytosol (**g**, n = 7) and mitochondria (**h**, n = 10) in the absence of glucose. The results of quantification in **e-h** are summarized in **d**. Error bar in **b-h** marks s.e.m.

An inhibitor of mitochondrial CiC, 1,2,3-benzenetricarboxylate (BTC), suppresses glucose-stimulated insulin secretion by blocking the mitochondrial export of citrate^8^. We found that the citrate concentration in INS-1 cells treated with 2 mM BTC after glucose addition exhibited a pronounced decrease from ~25 mM to ~10 mM, followed by an increase to ~18 mM (**Fig. 6d-f**). Cells treated with 2 mM BTC in the absence of glucose decreased in citrate concentration from ~3 mM to less than 2 mM (**Fig. 6d,g,h**). The citrate concentration in the cytosol and mitochondrial are found to be nearly identical and changed in parallel during our experiments. Imaging with the non-binding variant CitronRH confirmed that the fluorescence changes are indeed attributable to changes in citrate concentration (**Fig. S13**).

## Discussion

We have engineered two new high-performance citrate biosensors, Citron1 and Citroff1, through the use of semi-rational design and directed protein evolution. The initial prototype for both of these new biosensors was constructed by effectively inserting the CitA sensing domain into GFP at the uniquely permissive site in the vicinity of residues 146-147. Insertions at this site are generally well tolerated and such hybrid proteins (that is, GFP with a conformationally response domain inserted at this site) can generally be optimized to produce high performance constructs such as the GCaMP series of Ca^2+^ biosensors. In such biosensors, a conformational change in the sensing domain (for example, as induced by binding to an analyte of interest), is communicated to the GFP chromophore environment in such a way that the fluorescence intensity is reversible modulated.

To develop biosensors with effective coupling between the sensing domain and the GFP chromophore environment, we first attempted to optimize the two linkers that connect the CitA domain to GFP. This effort resulted in two distinct variants: one with a direct fluorescence response, and one with an inverse fluorescence response, to citrate binding. Further optimization by random mutagenesis and screening ultimately led to the Citron1 and Citroff1 biosensors, respectively. Variants of Citroff1 with *K*_d_ values ranging from 5 μM to ~1 mM were engineered by introducing rationally-designed mutations in the binding pocket or at previously reported positions^38^.

As a series, Citron1, Citroff1 (and its affinity variants) exhibit substantially larger dynamic responses, and much more sensitivity in the physiologically-relevant range of citrate concentrations, than the previously reported single FP-based citrate biosensors CF98. Furthermore, the availability of citrate-insensitive but pH-sensitive variants provides researchers with an ideal experimental control to confirm that signal changes from the biosensor are attributable to citrate rather than other physiological changes such as in pH. Together, these advantages mean that the advent of Citron1 and Citroff1 represent a major advance for the detection of citrate concentration dynamics in live mammalian cells using fluorescence imaging. Notably, the CitAP domain of the thermophilic bacterium *Geobacillus thermoleovorans* CitA SHK has recently been shown to be amenable to “binding pocket grafting” to convert its binding specificity to that of close homologs including L-malate, phthalate, and ethylmalonate^52^. Potentially, similar binding pocket grafting could be used to create a series of new biosensors based on the Citron1 and Citroff1 templates.

We have demonstrated the utility of these new citrate biosensors for the quantitative determination of citrate steady state concentrations in both the mitochondria and cytosol of HeLa and INS-1 cells. In addition, we have used these biosensors to follow dynamic changes in citrate concentration in cells induced by changes in glucose concentration in the media or by pharmacological treatment. These imaging results have generally reaffirmed what was already known, or expected, regarding citrate concentrations in these two cell types. For example, INS-1 cells were found to have substantially higher concentrations of citrate than HeLa cells, consistent with the higher metabolic rates of INS-1 cells^49^ However, these citrate indicators also enable intracellular citrate concentrations to be probed in ways that would be otherwise impractical or impossible. As one example, these indicators have enabled us to determine that, upon treatment of starved cells with glucose, the increase in cytosolic citrate concentration is delayed ~50 s relative to the increase in mitochondrial citrate concentration. As another example, these indicators have enabled us to observe transient changes in citrate concentration that would likely escape notice using traditional biochemical approaches that lack high temporal precision. For example, treatment of cells with the ACLY inhibitor BMS-303141 caused a transient increase in citrate concentration followed by a sustained decrease in concentration. In summary, we expect these high-performance citrate biosensors will see widespread adoption for use in tracking the metabolism of various types of cells for biochemical and pharmacological studies conducted either *in vitro* and *in vivo*. Both particularly low and particularly high concentrations of citrate have been reported to be key characteristics of normal and diseased cell types, and to play a role in a variety of fundamental processes. For example, citrate concentration in prostate cancer cells is substantially lower than in non-cancerous cells^53^, and astrocytes produce and release large amounts of citrate into cerebrospinal fluid^54^. In the extracellular milieu, citrate might function as a chelator of free Ca^2+^ and Zn^2+^, altering the excitable state of neurons^54^. To provide insight into this hypothesis, the green citrate biosensors reported here could potentially be used simultaneously for multi-parameter, multicolor imaging with red fluorescent Ca^2+^ and Zn^2+^ biosensors^32,39^ in order to correlate changes in citrate concentration with changes in the concentration of these divalent cations. Finally, it has been found that early onset epilepsy in children is related to loss-of-function mutations in Na^+^-coupled citrate transporter (SLC13A5) in neurons, but the mechanism remains unclear^55–58^. For these example biological problems, and many others not mentioned here, imaging citrate concentration changes using citrate biosensors will undoubtedly provide us with a deeper understanding of the underlying mechanism and further highlight citrate’s role as one of life’s most central metabolites.

## Methods

### Plasmid construction

A pBAD/HisB (Thermo Fisher Scientific) plasmid containing the gene for ncpGCAMP6 was used as the template and amplified by PCR using Q5 polymerase to prepare the vector containing ncpGFP^27^. A synthetic gene (gBlocks) for CitAP and overlap regions with the vector for assembly was purchased from Integrated DNA Technologies. The CitAP gene was cloned into the vector using Gibson assembly (New England Biolabs) and the resulting plasmid was used to transform DH10B electrocompetent cells. Plasmids were purified from 4 mL liquid culture of a single colony using the GeneJET miniprep kit and verified by sequencing.

### Linker optimization, directed evolution and *K*_d_ tuning

The two linkers were optimized separately using Quikchange II site-directed mutagenesis Lightening kit (Agilent) with customized primers. *E. coli* was transformed with gene libraries in the context of pBAD/HisB plasmids. Colonies showing high fluorescence intensity were picked and cultured in liquid media, followed by protein extraction with B-PER (Bacterial Protein Extraction Reagent, ThermoFisher Scientific), and fluorescence measurement in plate reader. A directed evolution strategy was employed to improve the dynamic range of the biosensors after linker optimization. Libraries were created using error-prone PCR as previously described^59^. In the first round of screening, ~25 colonies out of ~5000 colonies on agar plates were picked and cultured in 4 mL media using 15 mL culture tubes (Falcon). For the following rounds of directed evolution, 96 well culture blocks (Corning) were used for 1 mL liquid culture for higher-throughput screening (100 - 500 out of ~5000 colonies per round). Overnight cultures were spun down in centrifuge to pellet the bacteria, and B-PER was used to extract proteins for fluorescence measurement. Site-directed mutagenesis at single and double positions was utilized to change the citrate binding affinity. For screening purposes, each variant extracted from bacteria was tested in TBS buffer with 0.02, 0.2, 2 and 20 mM citrate.

### Protein purification and in vitro characterization

To purify the citrate biosensors for *in vitro* characterization, we first used the pBAD plasmids containing the gene encoding citrate biosensors to transform DH10B electrocompetent cells. Bacteria were then plated on LB-agar plates with 400 μg/mL ampicillin and 0.02% (wt/vol) arabinose for overnight incubation at 37 °C. The following day, colonies were picked and sub-cultured in 500 mL LB media containing 400 μg/mL ampicillin in a shaker at 37 °C for 24 h. The media was then supplemented with 0.02% (wt/vol) arabinose and shaken at room temperature for another 24 h. Bacterial cells were harvested using centrifuge and re-suspended in 50 mL Tris-buffered saline (TBS). The cells were lysed using sonication and the supernatant was obtained after centrifugation at 13,000 *g* for 30 min. The protein biosensors were purified by Ni-NTA agarose (MCLAB) and the TBS buffer was exchanged into MOPS buffer using 10 kDa ultrafiltration tubes (Amicon).

A Beckman DU-800 UV-vis spectrophotometer was used to measure absorption spectra and a Tecan Safire2 fluorescence plate reader was used to measure the excitation and emission fluorescence spectra. Extinction coefficient and quantum yield of each variant in the bound and unbound state was measured and calculated as previously described^60^. The brightness of the biosensor was determined as the product of extinction coefficient and quantum yield. To measure the binding affinity, proteins were diluted in various MOPS buffer containing citrate at the concentration ranging from 1 μM to 50 mM for fluorescence measurement. The fluorescence vs. concentration of citrate was plotted and fitted in Origin using Hill equation for determination of apparent *K*_d_. pH profiles of the proteins were similarly acquired. Proteins were diluted in a series of buffer with 30 mM sodium acetate and 30 mM borax (pH ranges from 4.5 to 11) for fluorescence measurement. Fluorescence vs. pH of the biosensor at bound and unbound state was plotted. For specificity test, the fluorescence spectra of each biosensor variant in MOPS buffer with or without 20 mM metabolite (citrate, malate, fumarate, succinate or α-ketoglutarate) were acquired.

### Crystallization and structure determination

The Citron1 recombinant protein was further purified with size exclusion chromatography using Superdex200 column (GE) in 20 mM Tris pH 8.0, 150 mM NaCl supplemented with 100 mM sodium citrate, and concentrated to 15 mg/mL for crystallization trials. The initial crystallization screen was carried out with 384-well plates (Molecular Dimensions) via sitting drop vapor diffusion against commercial sparse matrix screen kit (Hampton) at 20 °C. The recombinant Citron1 protein was crystallized in 4% v/v tacsimate pH 5.0, 12% w/v polyethylene glycol 3350. The Citron1 crystal was cryoprotected with the reservoir solution in supplement with 20% glycerol and flash frozen in liquid nitrogen. The dataset was collected in Shanghai Synchrotron Radiation Facility Beamline BL19U1 with wavelength of 0.98 Å and the X-ray diffraction data were processed and scaled with XDS package^61^. The data collection details and statistics are summarized in **Table S3**.

The Citron1 structure was solved with molecular replacement using both the GFP (6GEL) and CitA citrate-binding domain (2J80) as search model^25,62^. The density of the linker and citrate ions were observed after initial refinement, the manual model building of the missing residues and further refinement were carried out with COOT and PHENIX suite^63,64^. The Citron1 in complex with citrate was solved at resolution of 2.99 Å in the unit cell of a = b = 92.4 Å, c = 123.0 Å, and refined with a *R*_*work*_ of 0.2227 and *R*_*free*_ factor of 0.2822, respectively. The final Citron1 model contains two monomers and includes residues ranging from 2-361, 2 citrate ions and some residues from the cloning tag sequence. The structure exhibited good stereochemistry and showed a favorable Ramachandran plot. Detailed refinement statistics are provided in **Table S3**. Structure coordinates have been deposited in the Protein Data Bank with a code of 6LNP.

### HeLa cell imaging

The genes encoding the citrate biosensors were digested with XhoI and HindIII, and ligated into pcDNA3.1 (Thermo Fisher Scientific) vectors for cytoplasmic expression in mammalian cells. For mitochondria-targeting, genes of interest were amplified using Q5 PCR, purified, digested with BamH1 and HindIII, and ligated into a similarly-digested pcDNA3.1 vector with mitochondria-targeting sequence from cytochrome C oxidase signal peptide (MSVLTPLLLRGLTGSARRLPVPRAKIHSLGDP). Constructed plasmids were used to transform *E. coli* cells for plasmid amplification and purification with miniprep kits. HeLa cells were cultured in DMEM media with 10% fetal bovine serum and penicillin-streptomycin (Thermo Fisher Scientific). Cells were plated in either 35 mm glass bottom dishes or 24-well glass bottom plates, transfected using Turbofect (Thermo Fisher Scientific) according to the protocols provided in the manual, and incubated in a CO_2_ incubator for 24 - 48 h. Before imaging, cell culture media was removed and cells were washed twice with phosphate buffered saline (PBS), which was then replaced with HEPES buffered HBSS solution for imaging. Cell culture dishes or plates were fixed on the stage of a Nikon Eclipse Ti-E microscope or a Zeiss Axiovert 200 microscope with a 20✕ objective. Software was used to control the microscopes (NIS-Elements Advanced Research (Nikon) for the Nikon microscope and MetaMorph (Molecular Devices) for the Zeiss microscope). A GFP optical filter set (470/40 nm excitation and 525/50 nm emission) was used for imaging the citrate biosensors. The GFP filter set and an RFP optical filter set (535/50 nm excitation and 609/57 nm emission) was used and alternatively switched for dual-color imaging of the green citrate biosensors and the red ATP biosensor MalionR^43^.

For cell permeabilization experiments, time-lapse images were acquired every 10 or 20 s with appropriate exposure. Digitonin solution (0.1%) was added to reach a concentration of 0.0005% for plasma membrane permeabilization and 0.005% for mitochondrial membrane permeabilization after the imaging was initiated for ~100 s. A stock citrate solution (0.5 M) was then added to reach a concentration of 10 mM after the imaging was initiated for ~400 s. For *in situ* titration, cells were similarly permeabilized and various amounts of citrate solution were added to the buffer. After each citrate addition, cells were imaged for ~400 s to allow them to equilibrate and for the fluorescence to stabilize before the next addition.

To image pharmacologically-induced citrate concentration changes (with glucose, BMS-303141 or UK-5099), time-lapse images were acquired every 10 s to 10 min, with appropriate exposure times. For imaging of glucose treatment, cells were first starved in a glucose-free buffer for ~0.5 - 1 h before imaging. A solution of 500 mM glucose, 100 μM BMS-303141 or 1 mM UK-5099 was added to the imaging buffer to reach a final concentration of 20 mM for glucose, 5 μM for BMS-303141 or 25 μM for UK-5099, at about 2 min after the imaging was initiated. Due to the limited solubility in aqueous phase, UK-5099 was first dissolved in DMSO and then diluted with water. This resulted in a relatively high concentration of DMSO, a final concentration of ~0.25% (vol/vol) being added to the imaging buffer together with UK-5099. For control experiments, the same concentration of DMSO solution was used to treat HeLa cells expressing either Citron1 or CitronRH.

For dual color imaging of citrate and ATP concentration changes in mitochondria, the gene encoding MalionR (Addgene plasmid #113908) was amplified by PCR, digested with BamH1 and HindIII, and cloned into the mitochondria-targeting vector. An approximately 1:1 mixture of plasmids of mitochondria-targeted Citron1 and MalionR were used to co-transfect HeLa cells using the protocol described above. After 24 h incubation, cells were starved in a glucose free buffer for 0.5 - 1.0 h and then imaged. Glucose was added to induce citrate/ATP changes while imaging.

### INS-1 cell imaging

INS-1 cells were cultured in RPMI-1640 (Invitrogen) supplemented with 10% fetal bovine serum, 1% penicillin-streptomycin, 10 mM HEPES, 1 mM sodium pyruvate, 2 mM L-glutamine and 50 μM β-mercaptoethanol at 37 °C in a CO_2_ incubator. Cells were plated in 24-well glass bottom plates, and transfected using Lipofectamine 2000 (Invitrogen) according to the manual. Plasmids used for cytosolic and mitochondrial expression were the same as those used for HeLa cells. Before imaging, the culture medium was removed, INS-1 cells were washed twice with PBS buffer and Krebs-Ringer buffer (PH 7.4) was used for imaging. Krebs-Ringer buffer contains 0.1% BSA, 10 mM HEPES, 115 mM NaCl, 5 mM KCl, 24 mM NaHCO_3_, 2.5 mM CaCl_2_, and 1 mM MgCl_2_. Imaging was performed on a Nikon Eclipse Ti-E microscope or a Zeiss Axiovert 200M microscope. Citrate concentration in the cytosol or mitochondria was quantified similarly as HeLa cell experiments using *in situ* titration. To stimulate intracellular citrate changes, 20 mM glucose or 2 mM BTC was added to the imaging buffer and the corresponding citrate concentration changes were tracked with Citron1.

## Acknowledgements

The authors thank the University of Alberta Molecular Biology Services Unit for technical support, and Dr. Patrick E. McDonald and Dr. Wen-hong Li for helpful comments and providing INS-1 cells. MaLionR was a gift from Tetsuya Kitaguchi (Addgene plasmid # 113908). This work was supported by grants from the Natural Sciences and Engineering Research Council of Canada (NSERC; RGPIN 2018 04364), and the Canadian Institutes of Health Research (CIHR; FS 154310). YW was supported by the National Natural Science Foundation of China (NO.31870132, NO.81741088). We thank the staff from the BL19U1 beamline for technique support during data collection at the National Center for Protein Sciences Shanghai (NCPSS) at Shanghai Synchrotron Radiation Facility.

## Author contributions

Y.Z. developed Citron1 and Citroff1, performed protein characterizations, performed live cell imaging experiments, analyzed data, prepared figures, and wrote the manuscript. Y.W. crystallized the protein, solved the structure, and prepared figures. Y.S. performed protein characterizations, prepared figures, and wrote the manuscript. R.E.C. supervised research, prepared figures, and wrote the manuscript.

## Data availability

The data supporting this research are available upon request. Structure coordinates have been deposited in the Protein Data Bank with a code of 6LNP. Plasmid constructs encoding Citron1(#134300, #134303, #134305), Citroff1 (#134301, #134304), and other affinity variants (#134302, #134306, #134307) are available through Addgene.

## Supplementary Tables

**Table S1.**
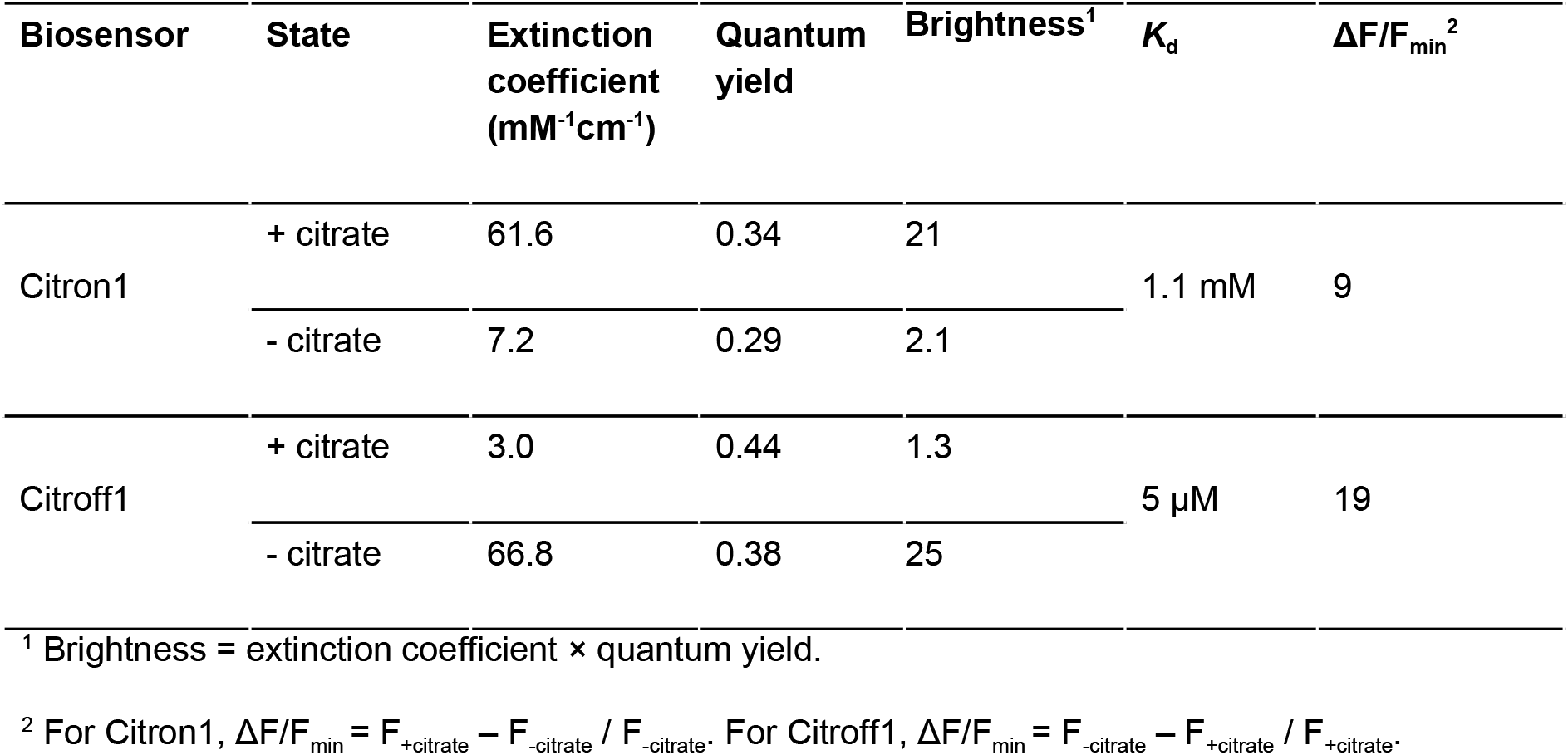
*In vitro* characterization of Citron1 and Citroff1.

| Biosensor | State | Extinction coefficient<br>(mM<sup>-1</sup>cm<sup>-1</sup>) | Quantum yield | Brightness<sup>1</sup> | $K_d$ | $\Delta F/F_{\min}$<sup>2</sup> |
| --- | --- | --- | --- | --- | --- | --- |
| Citron1 | + citrate | 61.6 | 0.34 | 21 | 1.1 mM | 9 |
|  | - citrate | 7.2 | 0.29 | 2.1 |  |  |
| Citroff1 | + citrate | 3.0 | 0.44 | 1.3 | 5 μM | 19 |
|  | - citrate | 66.8 | 0.38 | 25 |  |  |

<sup>1</sup> Brightness = extinction coefficient × quantum yield.
<sup>2</sup> For Citron1, $\Delta F/F_{\min} = F_{+\text{citrate}} - F_{-\text{citrate}} / F_{-\text{citrate}}$ . For Citroff1, $\Delta F/F_{\min} = F_{-\text{citrate}} - F_{+\text{citrate}} / F_{+\text{citrate}}$ .

**Table S2.**
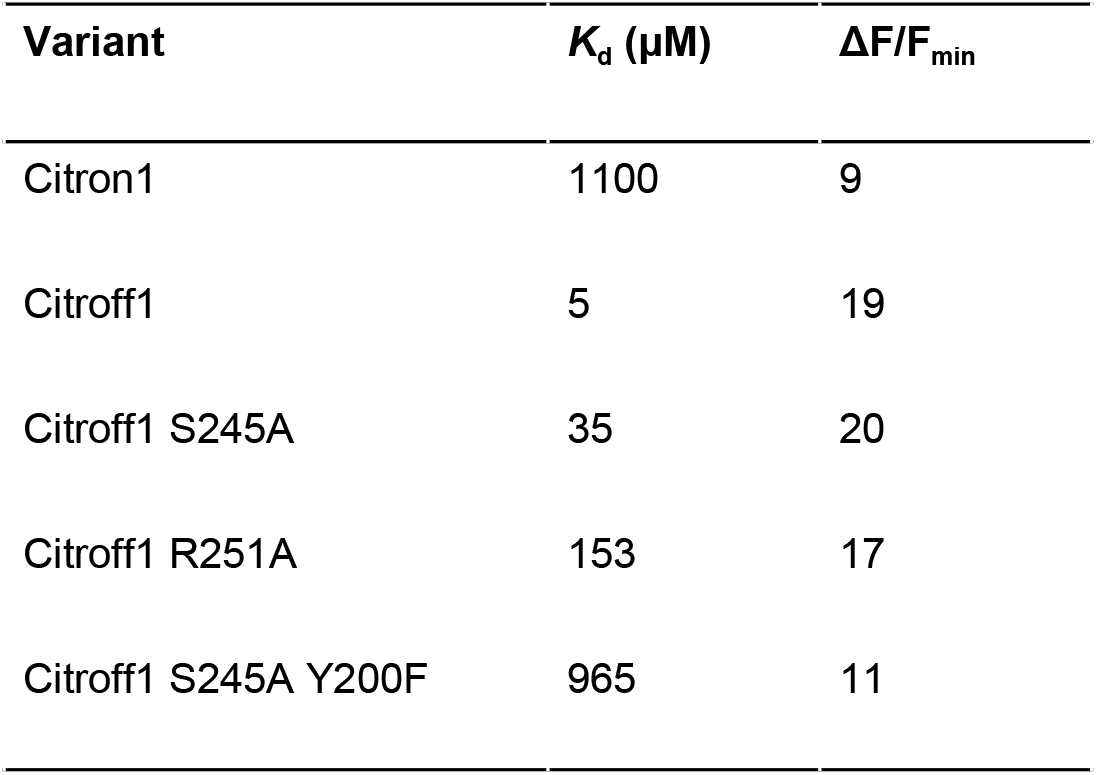
Summary of *K*_d_ and dynamic range of Citron1 and Citroff1 variants.

| Variant | $K_d$ ($\mu\text{M}$) | $\Delta F/F_{\min}$ |
| --- | --- | --- |
| Citron1 | 1100 | 9 |
| Citroff1 | 5 | 19 |
| Citroff1 S245A | 35 | 20 |
| Citroff1 R251A | 153 | 17 |
| Citroff1 S245A Y200F | 965 | 11 |

**Table S3.** X-ray data collection and refinement statistics. Statistics for the highest resolution shell are shown in parentheses.

|  |  |
| --- | --- |
| Crystal | Citron1 |
| **Data collection** |  |
| Spacegroup | P 41 |
| a, b, c (Å) | 92.4, 92.4, 123.0 |
| $\alpha$, $\beta$, $\gamma$ (°) | 90°, 90°, 90° |
| Resolution (Å) | 44.79-2.99 (3.10-2.99) |
| Rmerge | 0.134 (1.37) |
| Rmeas | 0.143 (1.46) |
| Multiplicity | 13.6 (14.2) |
| CC(1/2) | 0.998 (0.65) |
| CC* | 0.999 (0.887) |
| I/$\sigma$(I) | 12.1 (1.1) |
| Completeness (%) | 99.76 (99.86) |
| Wilson B-factor (Å<sup>2</sup>) | 52.55 |
| **Refinement** |  |
| Total Reflections | 284653 (29409) |
| Unique Reflections | 20881 (2072) |
| $R_{\text{work}}/R_{\text{free}}$ | 0.2227/0.2822 |
| **Number of atoms:** |  |
| Protein | 5518 |
| Ligands | 70 |
| Protein Residues | 733 |
| Average B-factor (Å<sup>2</sup>) | 51.33 |
| Protein ADP (Å<sup>2</sup>) | 51.29 |
| Ligands (Å<sup>2</sup>) | 54.52 |
| **Ramachandran plot:** |  |
| Favored/Allowed (%) | 92.1/7.5 |
| **Root-Mean-Square-Deviation:** |  |
| Bond lengths (Å) | 0.004 |
| Bond Angle (°) | 0.80 |
| PDB code | 6LNP |

## Supplementary Figures

**Figure S1.**
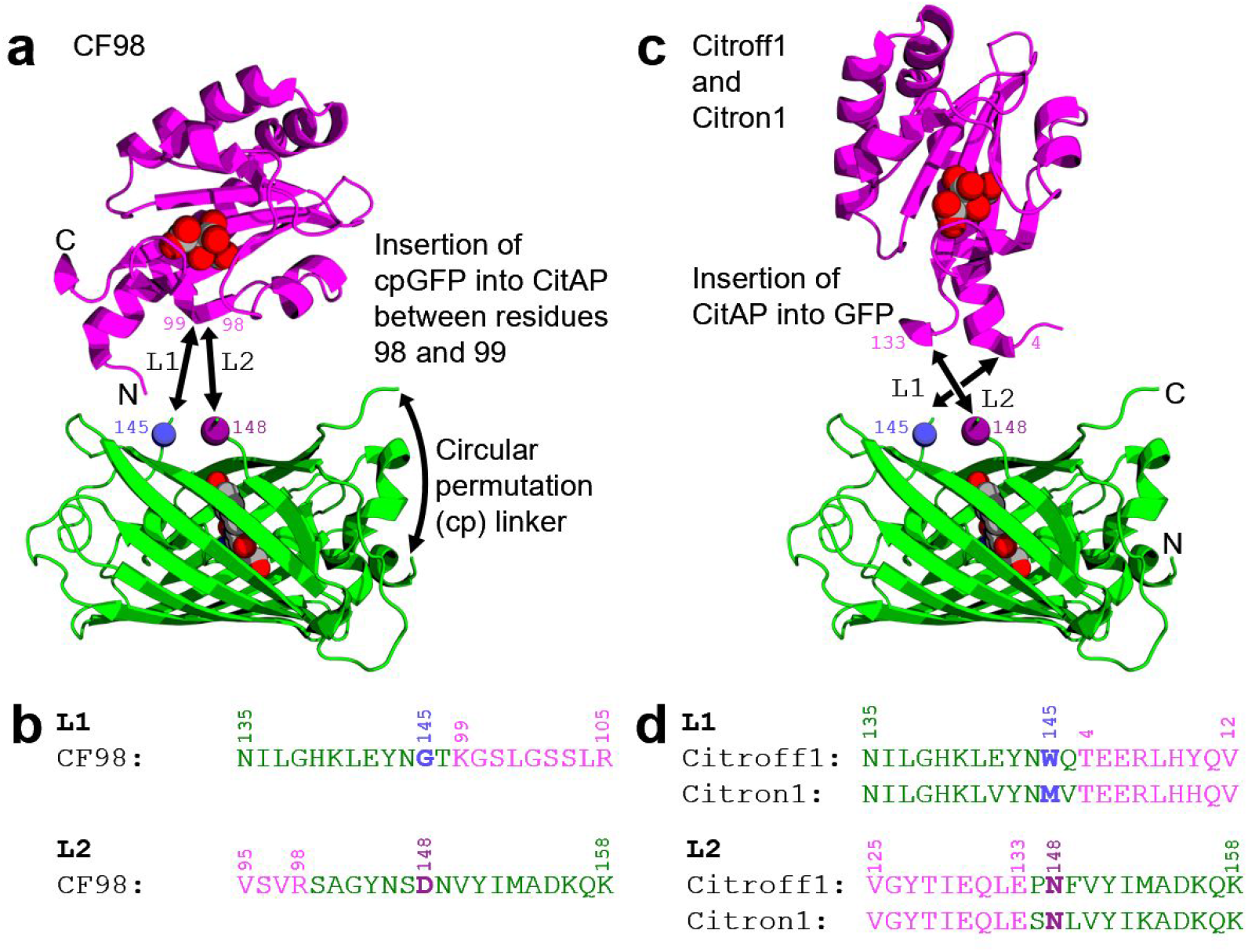
Comparison of the design of the previously reported CF98 biosensor and the design of the Citroff1 and Citron1 biosensors described in this work. Residues are numbered according to the original CitAP and GFP sequences. **a** CF98 was designed based on the rationale of propagating the binding-induced conformational change in the minor loop of CitAP (refer to **Fig. 1b**) into a change in the GFP chromophore environment^28^. To realize this design, cpGFP was inserted into the minor loop between residues 98 and 99. The CitAP domain with bound citrate is shown in magenta (PDB ID 2J80)^25^ and the GFP domain from GCaMP6m is shown in green (PDB ID 3WLD)^65^. Residues 145 (Cα represented as blue sphere) and 148 (Cα represented as a purple sphere) are shown as reference points. Black arrows are used to represent the linkage from GFP to residue 99 of CitAP (L1), the linkage from residue 98 of CitAP to GFP (L2), and the cp linker that connects the original N- and C-termini (which was disordered and not observed in this structure). **b** Sequences in the vicinity of the L1 and L2 connections between GFP and CitAP in the CF98 biosensor. Residues 145 and 148 are colored as in **a**. **c** In this work we describe citrate biosensors based on the rationale of propagating the binding induced conformational changes at the C-terminus of CitAP (refer to **Fig. 1**) into a change in the GFP chromophore environment. To realize this design, the CitAP domain was inserted into GFP at the same site where CaM-RS20 is inserted in the ncpGCaMP6s^27^ biosensor (a topological variant of GCaMP6s)^36^. Protein structures are represented as in **a**. L1 is the connection between GFP and residue 4 of CitAP, and L2 is the connection between residue 133 of CitAP and GFP, and there is no cp linker in this design. **d** Sequences in the vicinity of L1 and L2 for the Citroff1 and Citron1 biosensors. Residues 145 and 148 are colored as in **c**.

**Figure S2.**
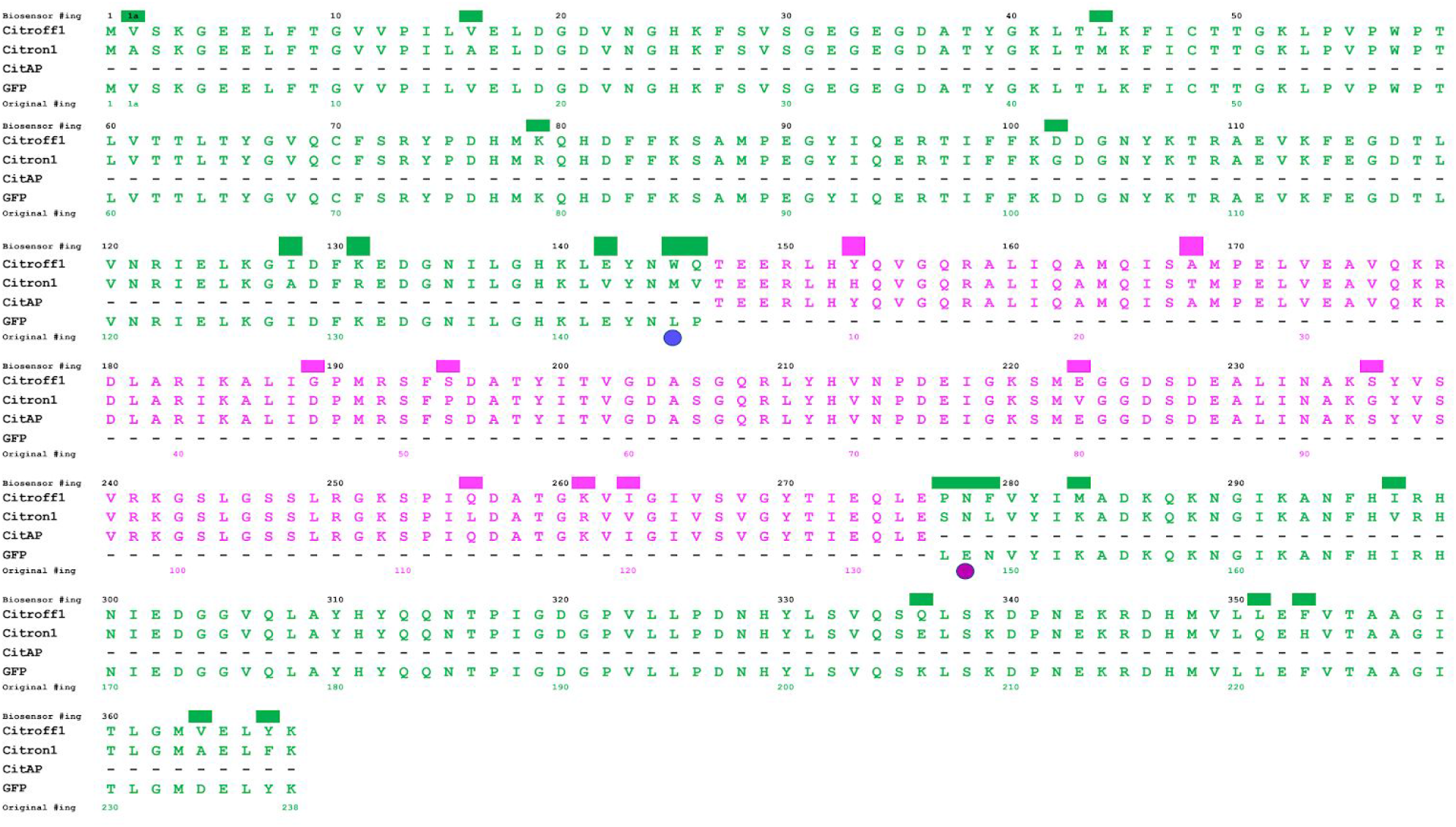
Sequence alignment of citrate biosensors described in this work. The CitAP sequence (magenta) is from, and numbered (bottom) as in, PDB ID 2J80^25^. The GFP sequence (green) is from ncpGCaMPs^27^ and numbered (bottom) according to wild-type GFP^66^. Numbers at the top correspond to residue numbering for the full length biosensors. The positions of mutations accumulated during biosensor optimization are represented with colored bars over the position of the mutation. Relative to the starting template sequences, Citroff1 acquired 5 mutations in linker regions, 3 mutations in GFP, and 1 mutation in CitA. Citron1 acquired 5 mutations in linker regions, 13 mutations in GFP, and 8 mutations in CitA.

**Figure S3.**
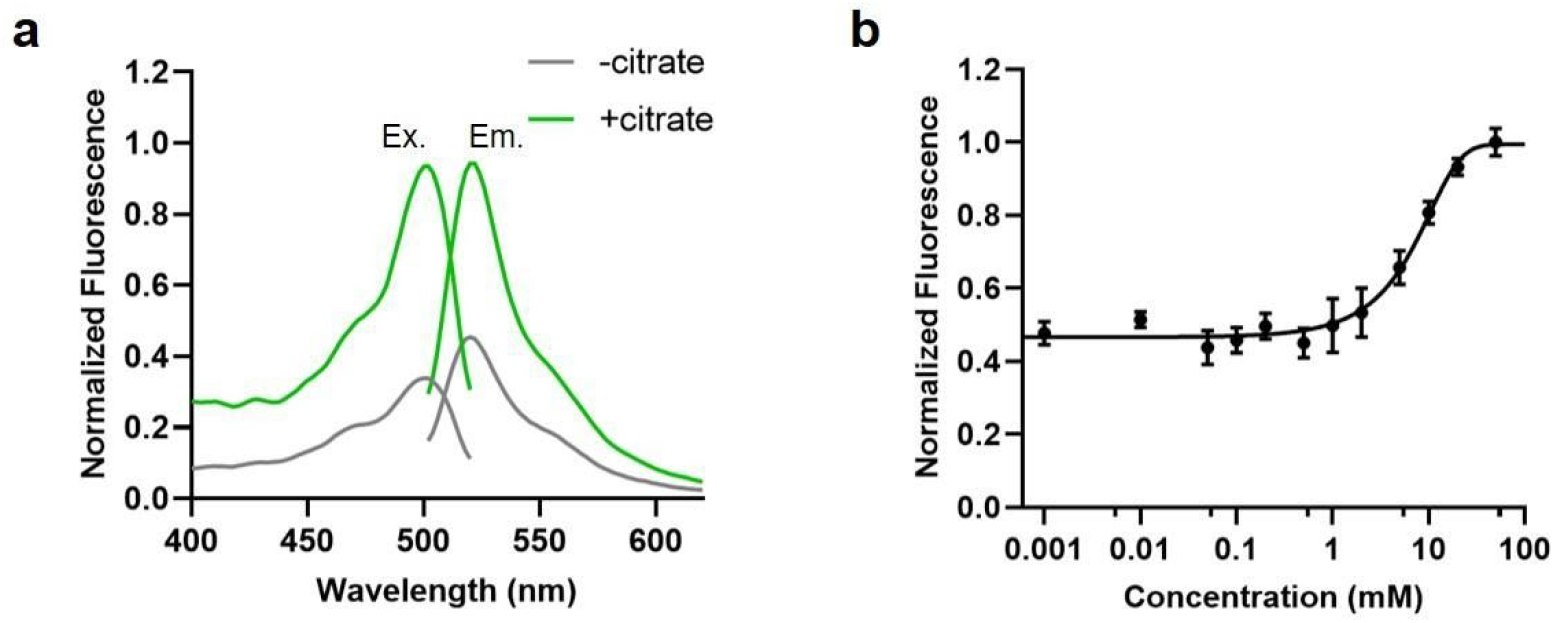
*In vitro* characterization of CF98. **a** Normalized excitation and emission spectra of purified CF98. **b** *In vitro* citrate titration curve of purified CF98. Error bars represent the standard deviation of triplicates.

**Figure S4.**
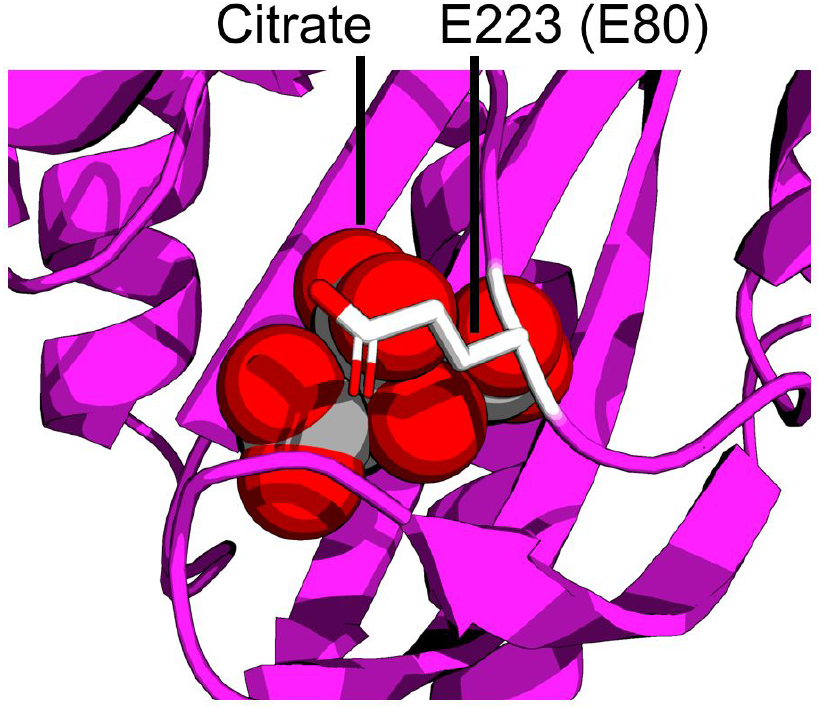
Location of the E223V mutation (equivalent to E80V as numbered in PDB ID 2J80)^25^ in Citron1.

**Figure S5.**
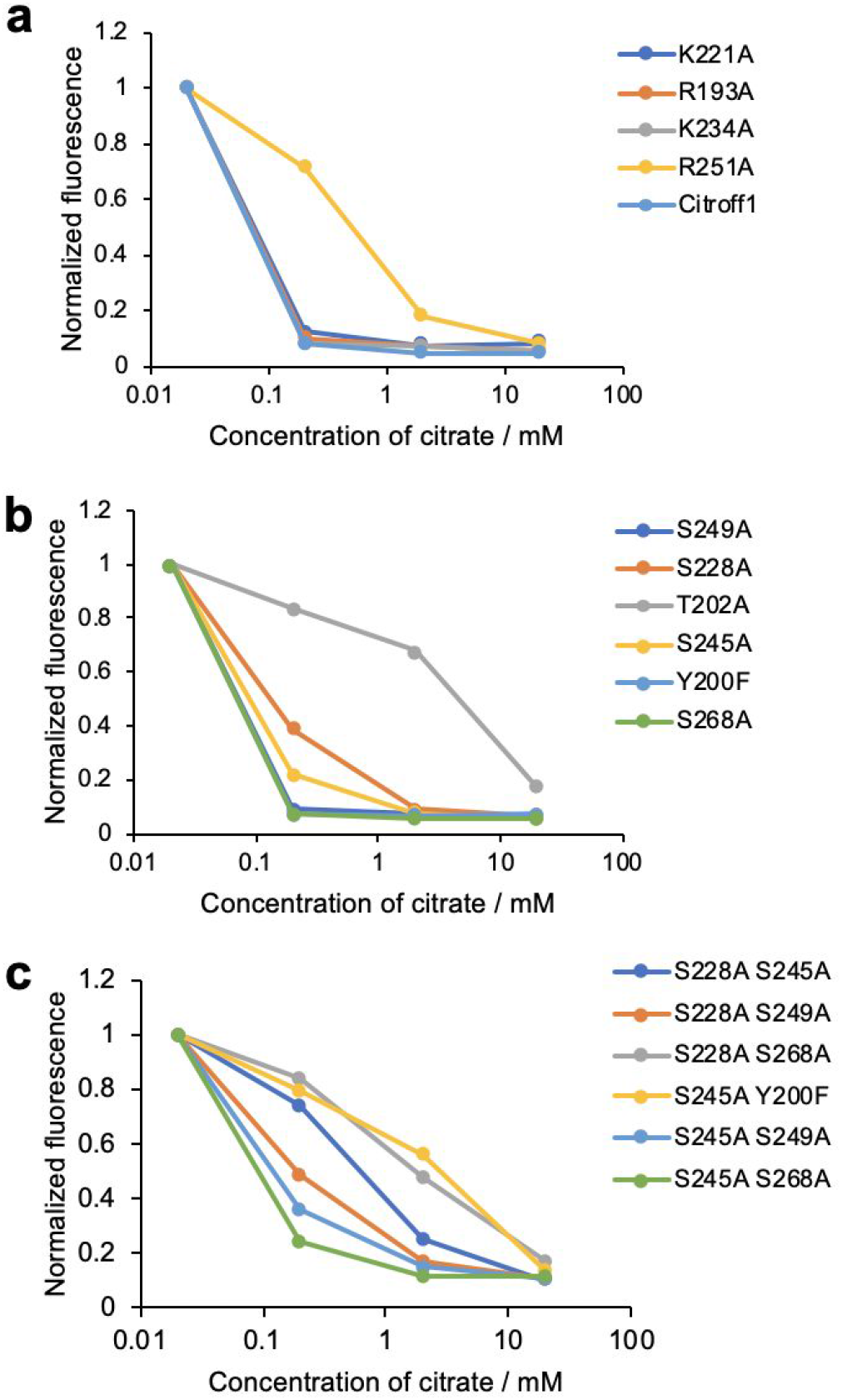
Introduction of mutations to decrease the binding affinity of Citroff1. **a** Introduction of mutations that were previously reported to decrease the affinity of a CitAP-based biosensor^28^. **b** Introduction of various structure-guided mutations that were hypothesized to potentially decrease the affinity for citrate. **c** Pairwise combinations of mutations.

**Figure S6.**
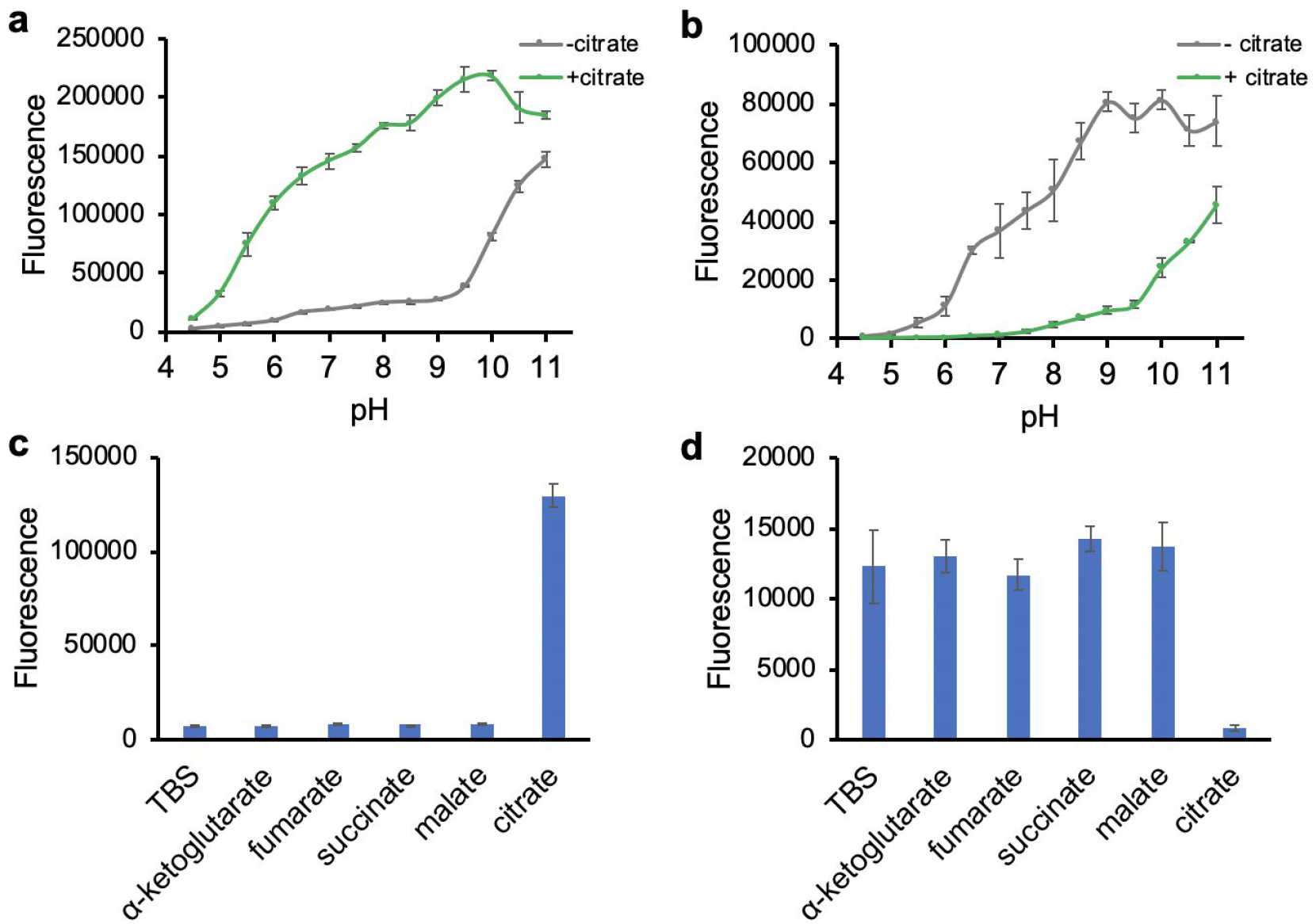
pH dependence and specificity of citrate biosensors. **a,b** Fluorescence response of Citron1 (**a**) and Citroff1 (**b**) at various pH values with or without 20 mM Citrate. **c,d** Specificity of Citron1 (**c**) and Citroff1 (**d**) to various metabolites (20 mM each metabolite in TBS buffer). Error bars represent the standard deviation of triplicates.

**Figure S7.**
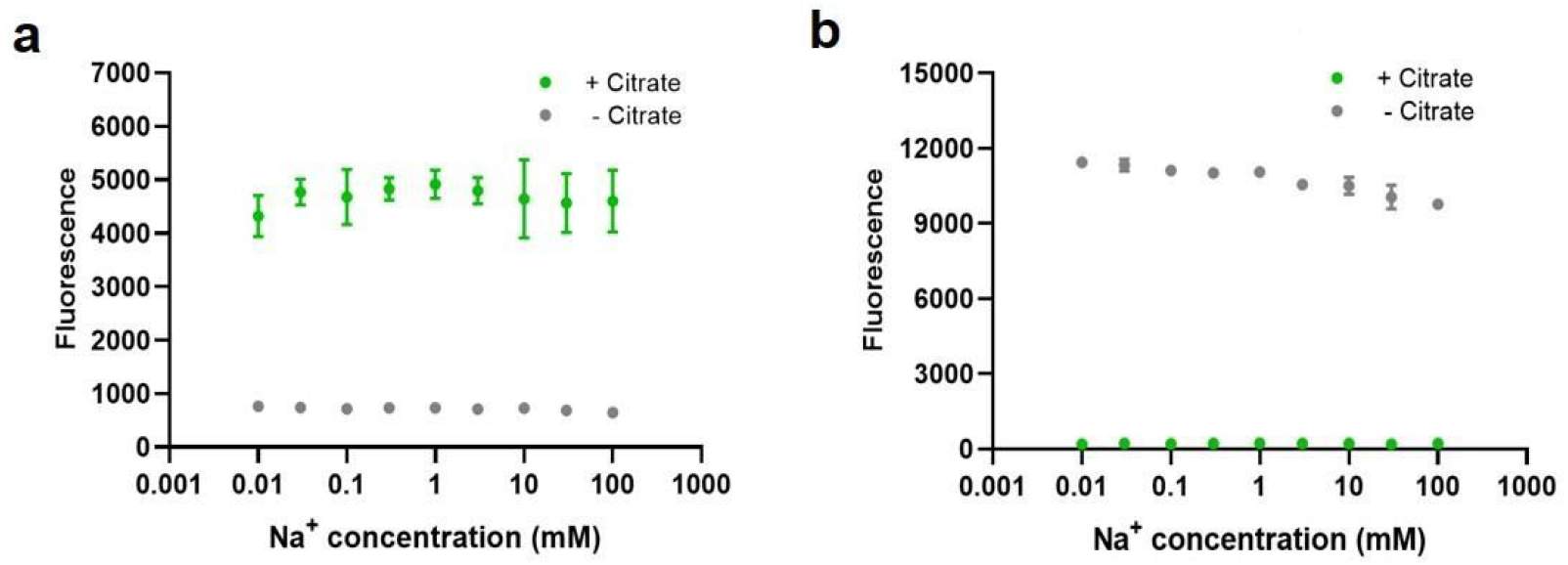
Na^+^ concentration dependence of Citron1 (**a**) and Citroff1 (**b**), with 30 mM citrate in the + citrate condition. Error bars represent the standard deviation of triplicates.

**Figure S8.**
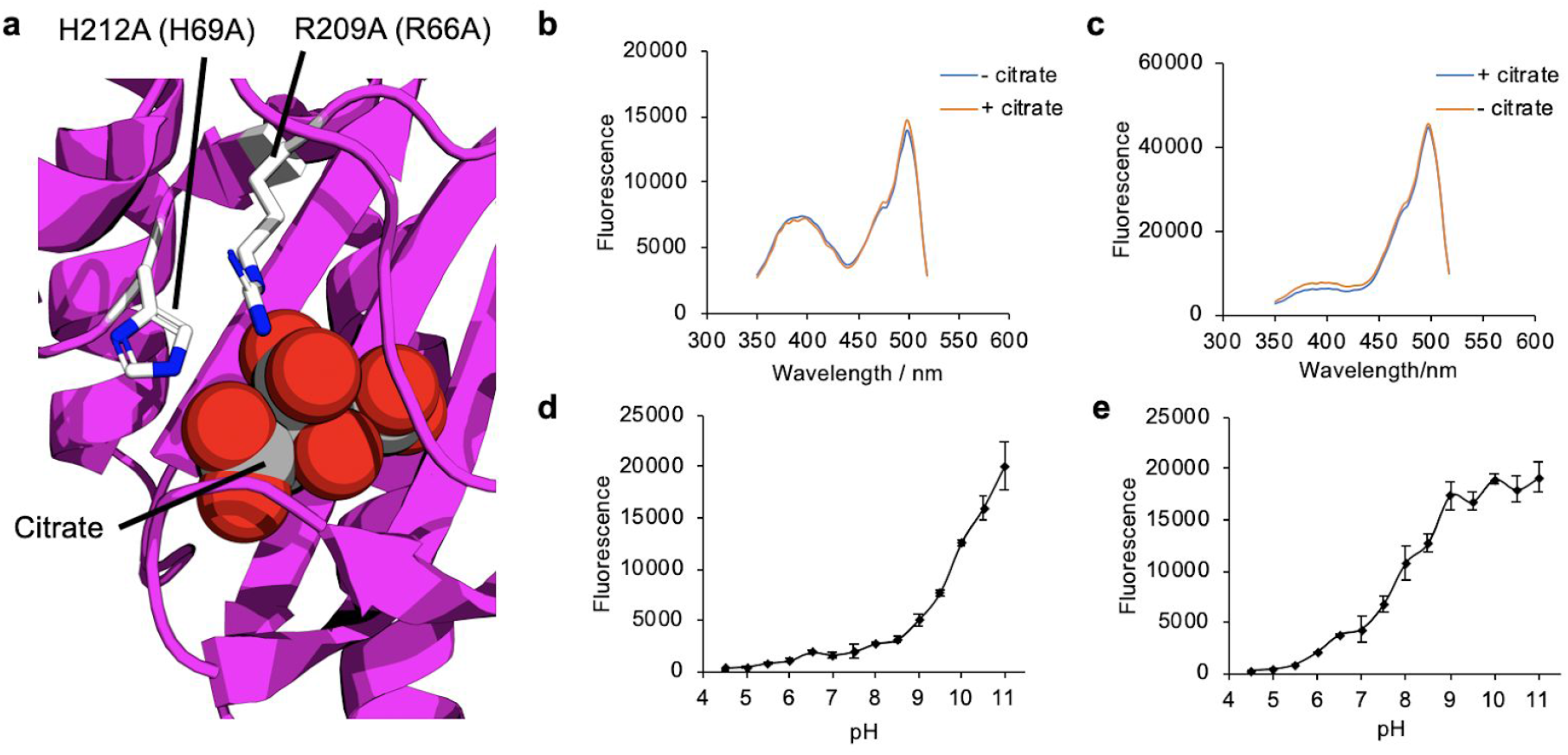
Engineering citrate-insensitive control constructs. **a** Crystal structure of CitAP (PDB ID 2J80)^25^ with side chains of H212 and R209 (biosensor numbering, equivalent to H69 and R66 of CitAP) shown in stick representation. Mutation of these two residues to alanine was sufficient to disable citrate binding. **b,c** Excitation spectra of control variants CitronRH (**b**) and CitroffRH (**a**) with or without citrate in the buffer. **d,e** pH profiles of CitronRH (**d**) and CitroffRH (**e**). Error bars represent the standard deviation of triplicates.

**Figure S9.**
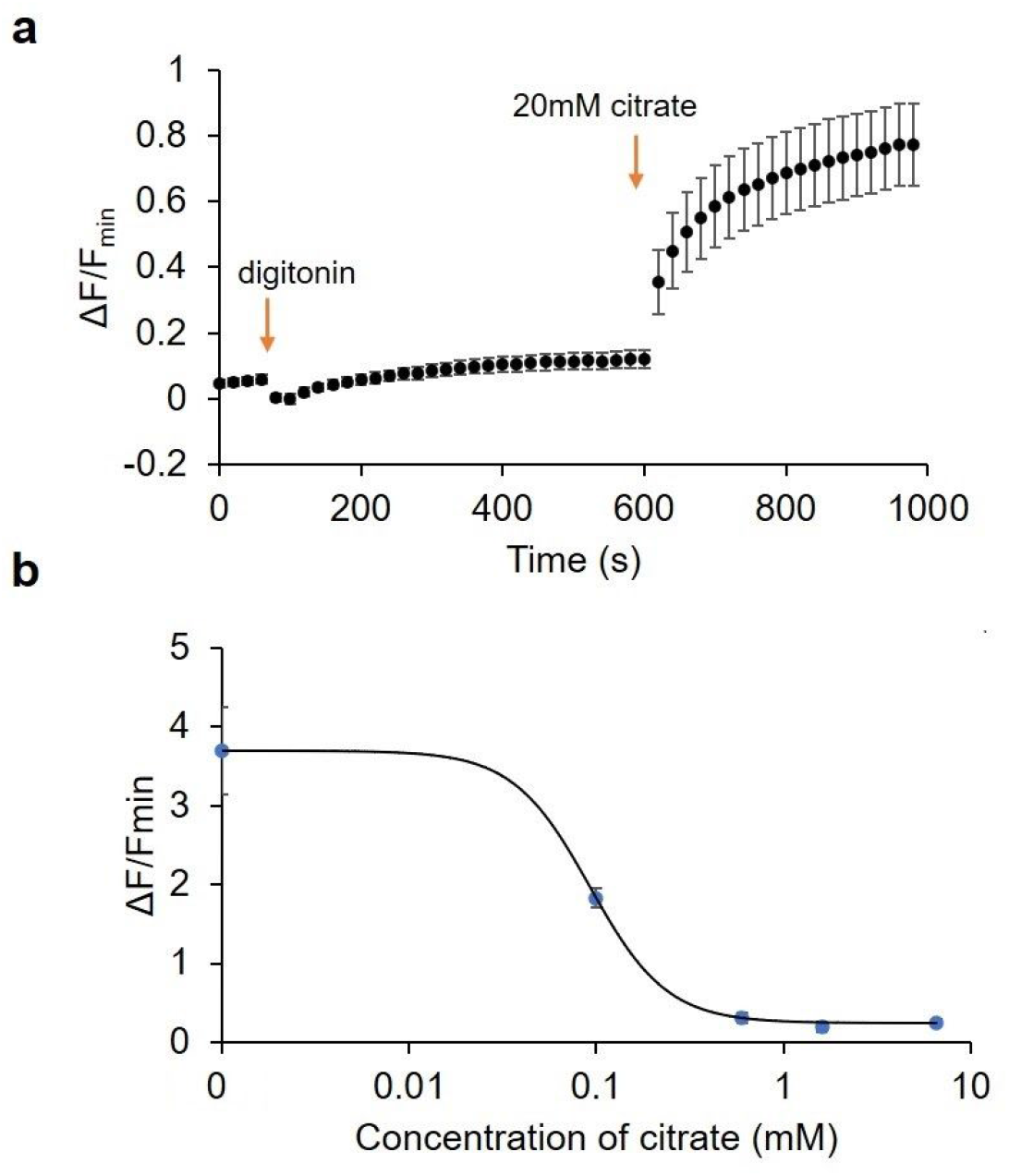
Additional information related to *in situ* calibration of citrate biosensors. **a** Citron1 expressed in the mitochondria of HeLa cells in response to digitonin and citrate (n = 22). **b***In situ* titration curve of Citroff1 in the cytosol (n = 26). Error bars represent s.e.m..

**Figure S10.**
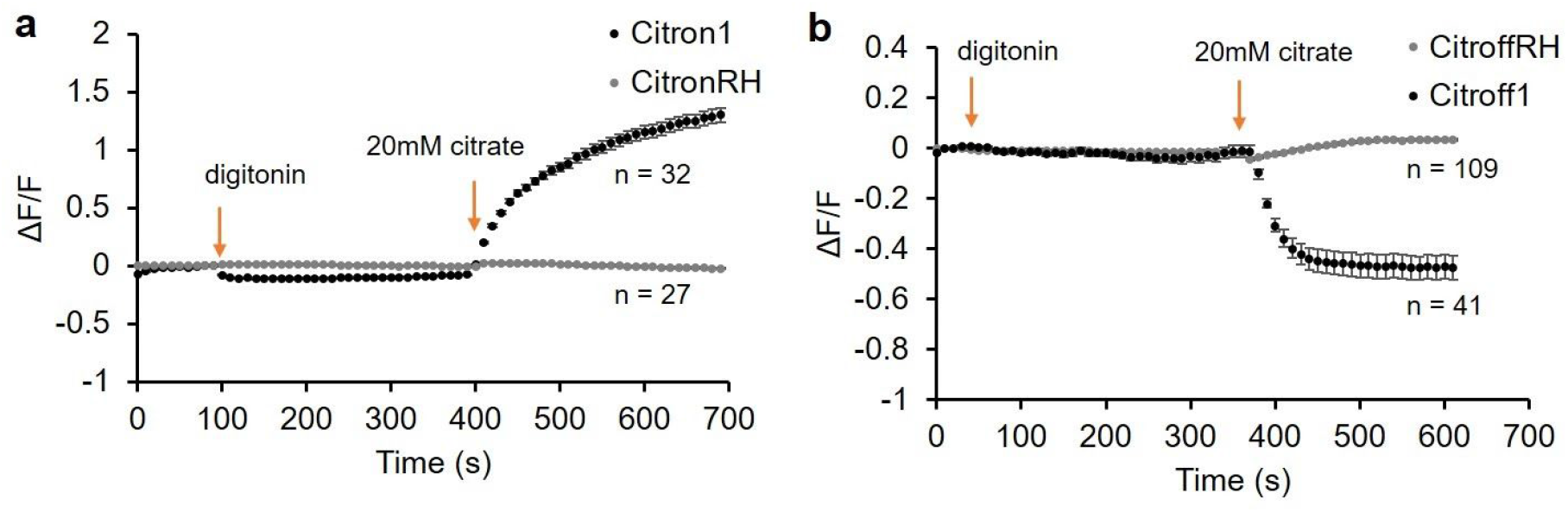
Comparison of citrate biosensors (Citron1 and Citroff1) and control variants (CitronRH and CitroffRH) expressed in the cytosol. Cells were treated with digitonin and citrate. Error bars represent s.e.m..

**Figure S11.**
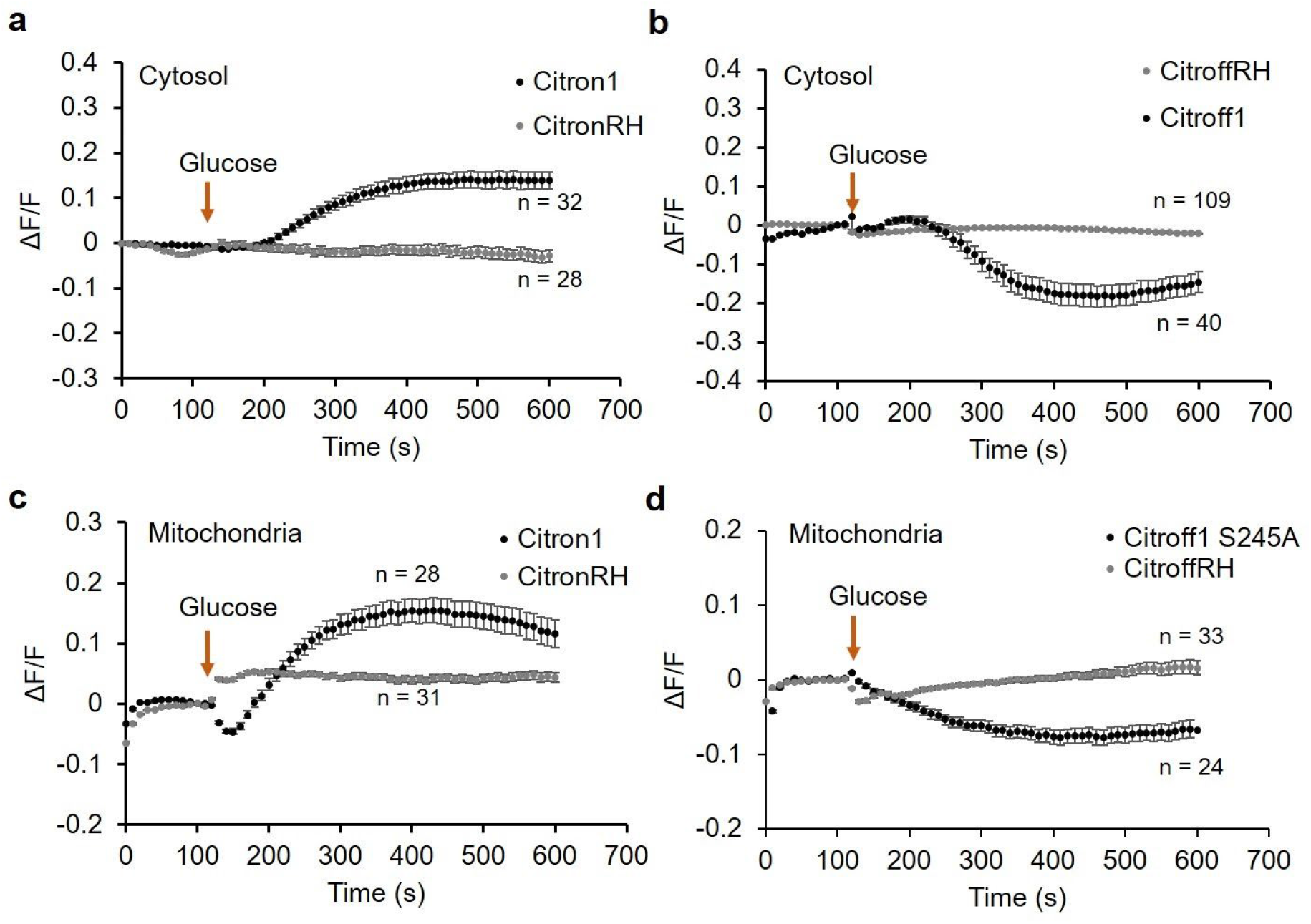
Responses of citrate biosensors (Citron1 and Citroff1 S245A) and control variants (CitronRH and CitroffRH) expressed in the cytosol (**a**,**b**) and mitochondria (**c**,**d**) of HeLa cells during treatment with glucose. Error bars represent s.e.m..

**Figure S12.**
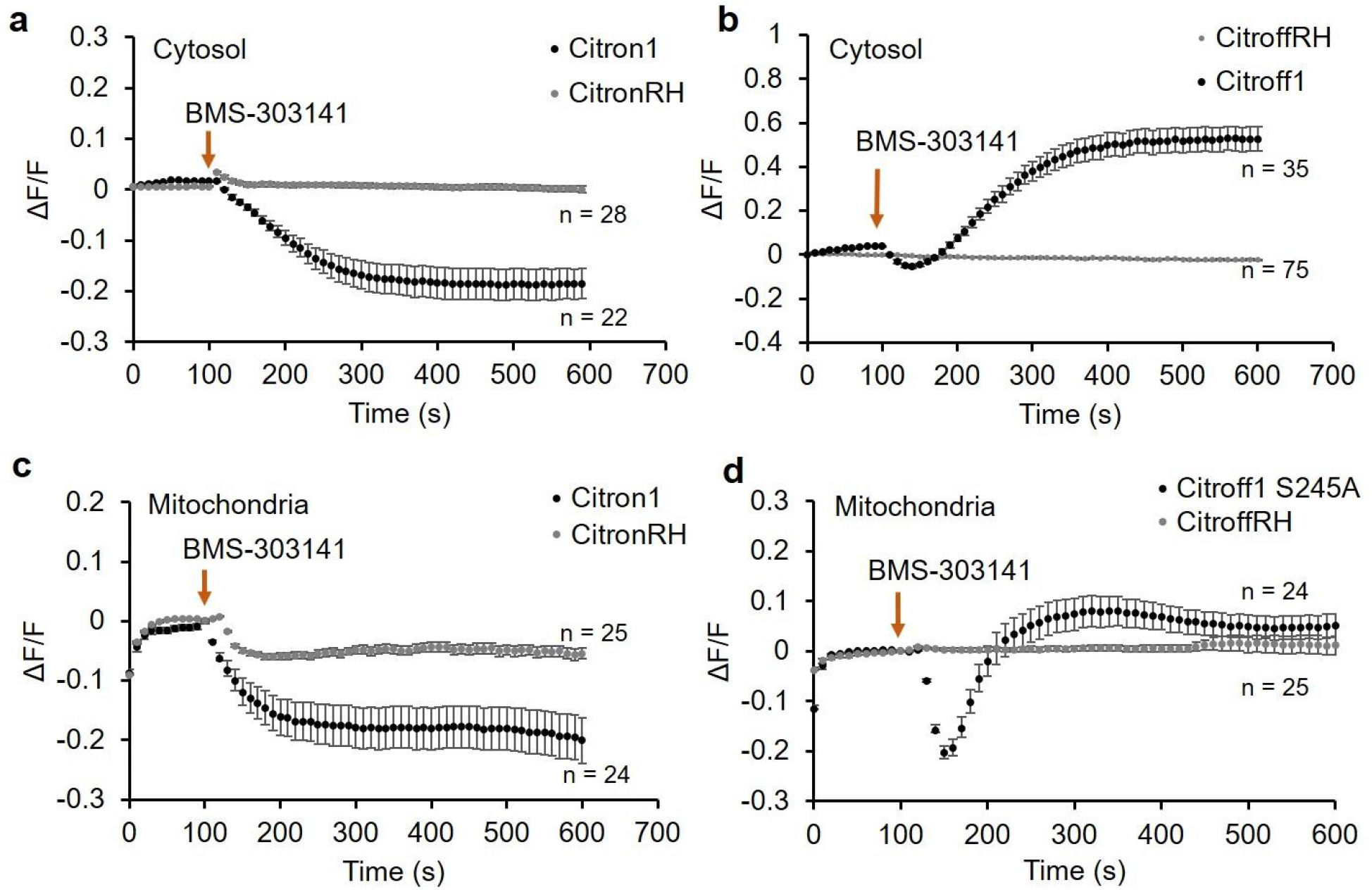
Comparison of citrate biosensors (Citron1 and Citroff1 S245A) and control variants (CitronRH and CitroffRH) expressed in the cytosol (a,b) and mitochondria (c,d) during treatment with BMS-303141. Error bars represent s.e.m..

**Figure S13.**
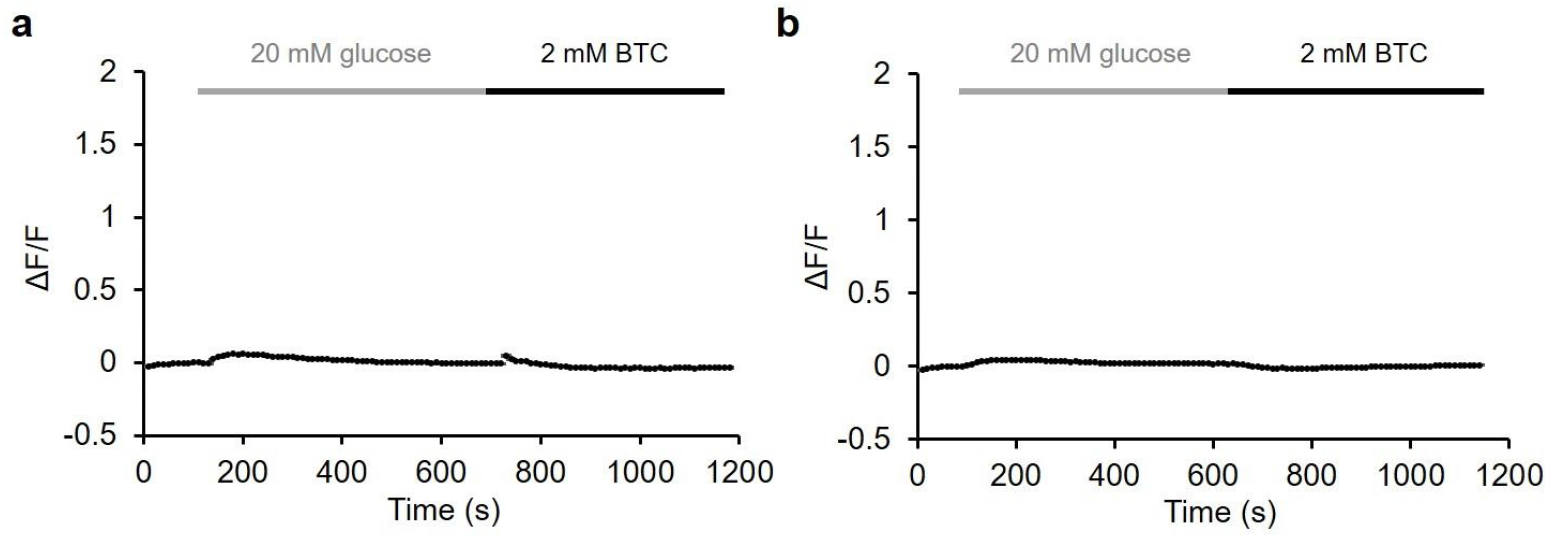
Imaging of the non-binding CitronRH variant in the cytosol (**a**, n = 23) and mitochondria (**b**, n = 16) of INS-1 cells. Cells were treated with 20 mM glucose and 2 mM BTC as indicated. Y-axis range is scaled to match **Fig. 6e**. Error bars represent s.e.m..

